# Deciphering stiffness-driven changes in colorectal cancer by proteomics

**DOI:** 10.1101/2024.10.16.618701

**Authors:** Charlotte Cresens, Ana Montero-Calle, Guillermo Solís-Fernández, Behrad Shaghaghi, Lotte Gerrits, Samet Aytekin, Paul H. J. Kouwer, Rodrigo Barderas, Susana Rocha

## Abstract

Tumor stiffening plays a pivotal role in cancer progression. Increased tumor stiffness, resulting from interactions between cancer cells and their surrounding microenvironment, alters the tumor’s mechanical properties and significantly impacts cancer growth and metastasis, the primary cause of cancer-related deaths. Despite the importance of tumor stiffness, systematic studies exploring its effect on proteomic profiles are limited. In this study, focused on colorectal cancer, we show that matrix stiffness significantly alters the expression of secreted proteins, while intracellular protein levels remain largely unaffected. Functional assays reveal that the changes in the secretome, driven by matrix stiffness, enhance cell migration, angiogenesis, and matrix remodeling, which collectively contribute to a more aggressive cancer phenotype. Our findings emphasize the critical role of matrix stiffness in driving colorectal cancer progression through changes in the secretome, offering valuable insights for the development of biomechanical cancer therapies.

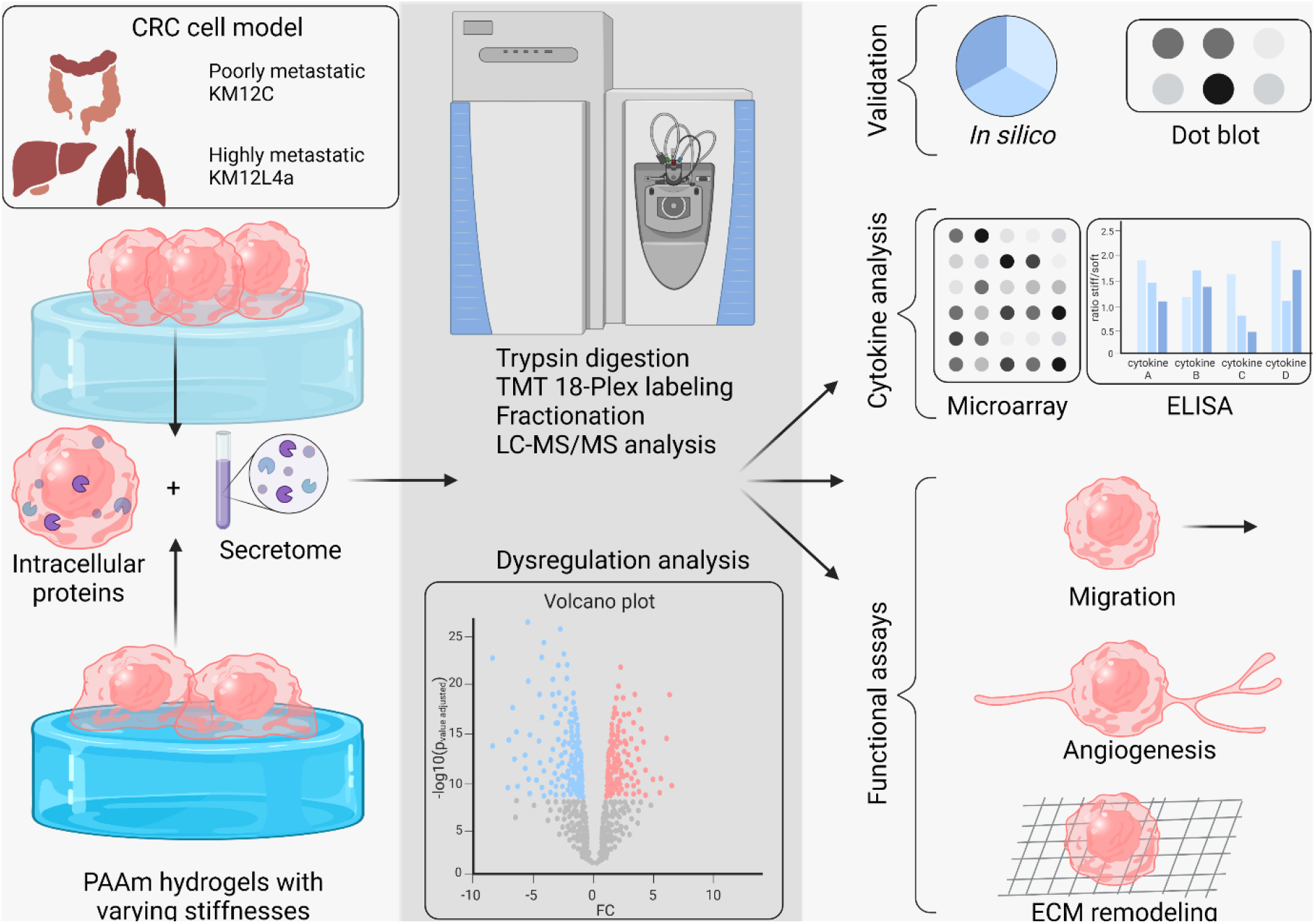

## Introduction

Colorectal cancer (CRC) ranks as the second most prevalent cancer in Europe when considering incidence rates across both genders.^1–3^ Predominantly sporadic, less than 5% of the cases occur in individuals with a hereditary predisposition. Despite the reduction in mortality rates over the past decade, owing to screening programs and advancements in treatments for both adjuvant and metastatic disease, CRC remains the second leading cause of cancer-related deaths, surpassed only by lung cancer.^4^ Metastasis, the process by which cancer cells spread from the primary site to other parts of the body, significantly contributes to CRC mortality. About one-third of CRC patients are diagnosed at an advanced stage, typically when the tumor has become stiffer and metastasis has already occurred. Since most patients with metastatic CRC require second- and third-line treatments as the disease progresses, the 5-year survival rate remains less than 10%. This high mortality rate underscores the importance of understanding and combating CRC metastasis, making the development of new approaches to study CRC metastasis essential.

Previous research has shown that matrix stiffness, a mechanical property of the extracellular matrix (ECM), plays a significant role in cancer progression, including CRC metastasis.^5^ Depending on the cancer type and stage of tumor progression, cancerous tissue can be up to 20 times stiffer than its healthy counterpart.^6^ For example, healthy colorectal tissue has a stiffness of ≈ 1 kPa, but as CRC progresses, stiffness typically increases. Early-stage tumors (T1 and T2) exhibit values ranging from 1 to 12 kPa, while more advanced stages (T3 and T4) and tumors with metastases show values between 3 and 68 kPa.^7^ This increase in stiffness is mainly driven by reciprocal interactions between cancer cells and other types of cells within the tumor microenvironment (TME), particularly cancer-associated fibroblasts (CAFs). These interactions promote the secretion of ECM proteins and enhance the activity of ECM-modifying enzymes (e.g. MMPs, ADAMs, TIMPs, LOX, and transglutaminases). Additionally, cellular traction forces further stiffen the tumor by aligning ECM fibers.^8–11^ The stiffened ECM has profound effects on cancer progression and metastasis, promoting the proliferation of tumor-initiating cells, epithelial-mesenchymal transition, alterations in extracellular vesicles, and various biochemical and biophysical changes in the TME.^12–16^

Despite its recognized importance, no study has yet systematically explored the effect of substrate stiffness at the protein level across the whole secretome, with previous work restricted to cell extracts or extracellular vesicles and outside the context of CRC.^17–21^ In this study, we examine the effect of increased substrate stiffness on CRC progression through proteomic-based analyses. We investigate this using the KM12 cell model of CRC metastasis, particularly the KM12L4a cell line, known for its high metastatic potential to liver and lung in nude mice, and its poorly metastatic isogenic counterpart, KM12C.^22,23^ The KM12 model successfully replicates key aspects of CRC metastasis, and comprehensive analysis of its transcriptome and proteome have provided valuable insights into the metastatic process.^24–32^ To explore stiffness-induced changes, we grow these cells on synthetic polyacrylamide (PAAm) hydrogels with varying stiffnesses, representative of different stages of CRC development and progression. We analyze intracellular and secreted proteins using mass-spectrometry based quantitative proteomics, known for its ability to identify and quantify thousands of proteins with high specificity and sensitivity. This enables a comprehensive analysis of protein expression, functionality, and interactions and is therefore a powerful tool for unraveling the complexities of biological systems. We validate our findings through dot blots, and extend our analyses to human cytokines by antibody microarrays and ELISAs. Finally, we perform various functional assays to evaluate the effects of substrate stiffness on cell behavior. This study underscores the significant role of ECM stiffness in CRC development and progression. Our findings highlight that understanding these processes and targeting elements of the physical tumor microenvironment, including matrix stiffness, could be key to overcoming drug resistance and unlocking new avenues for therapeutic development.^33^

## Materials and methods

### PAAm hydrogels

#### Hydrogel synthesis

Polyacrylamide (PAAm) hydrogels of various stiffnesses were synthesized as large, unattached gels as previously reported.^34,35^ Briefly, specified quantities (Table S1) of acrylamide (AAm, Sigma-Aldrich, Merck, A4058) and N,N’-methylenebisacrylamide (MBAA, Sigma-Aldrich, Merck, M1533) were mixed with distilled water (dH_2_O) and sonicated on ice for 10 min. Ammonium persulfate (APS, Bio-Rad, #1610700EDU) and N,N,N’,N′-tetramethylethylenediamine (TEMED, Sigma-Aldrich, Merck, T7024) were added and 9.5 mL of the mixture was polymerized at room temperature in a Petri dish (92 × 16 mm, Sarstedt, 82.1473.001). The lid served as the bottom support, and the base dish with open side up as the upper support. The resulting hydrogel, not attached to any surface and approximately 1.2 - 1.5 mm thick, was stamped either into one circular gel with 75 mm diameter for large-batch cell culture applications, or into multiple circular gels with 20 mm diameter for rheological characterization. Care was taken to avoid the outer 0.5 cm of the polymerized gel where oxygen can enter, since this is known to inhibit the polymerization and therefore could have affected the local mechanical properties.^34^ The stamped gel was washed three times with 10 mL dH_2_O, and then submerged in 100 mL DPBS (1x, prepared from Gibco, Thermo Fisher Scientific, 14200075) in a humidified incubator at 37 °C overnight for swelling.

#### Hydrogel characterization

PAAm hydrogels were mechanically characterized according to previously established protocols.^34,35^ Measurements were conducted using a stress-controlled rotational rheometer (TA Instruments, Discovery HR-2) equipped with a 20 mm parallel plate geometry (DHR & AR-G2, Stainless Steel, 20 mm Plate, NST, 511200.906). To prevent slipping on the already polymerized gel, sanding paper was attached to the geometry and lower plate, both hydrated with 1 mL DPBS to prevent hydrogel dehydration by the sanding paper. A swollen hydrogel was restamped into a 20 mm circular gel to match the geometry’s dimensions, and meticulously positioned on the rheometer to align with the sanding paper. The gel was compressed 3%, achieved by reducing the gap distance 3% for a gel with uniform geometry contact and non-negligible axial force of minimally 0.01 N. Then, a 30-second time sweep measurement was performed at 21 °C with 1% strain and 1 Hz frequency. Hydrogel stiffness (Youngs’ modulus, E) was calculated from the storage modulus (*G′*, taken as the average value across the time sweep) using the formula: E ≈ 2*G′* (1 + v), where v denotes the Poisson’s ratio (approximately 0.5 for PAAm gels).^36,37^ Average PAAm stiffness values, along with standard errors [average^*^(standard deviation/maximum)] were calculated from at least three technical replicates from the same large gel (Table 1, Table S1 for full dataset).

**Table 1:**
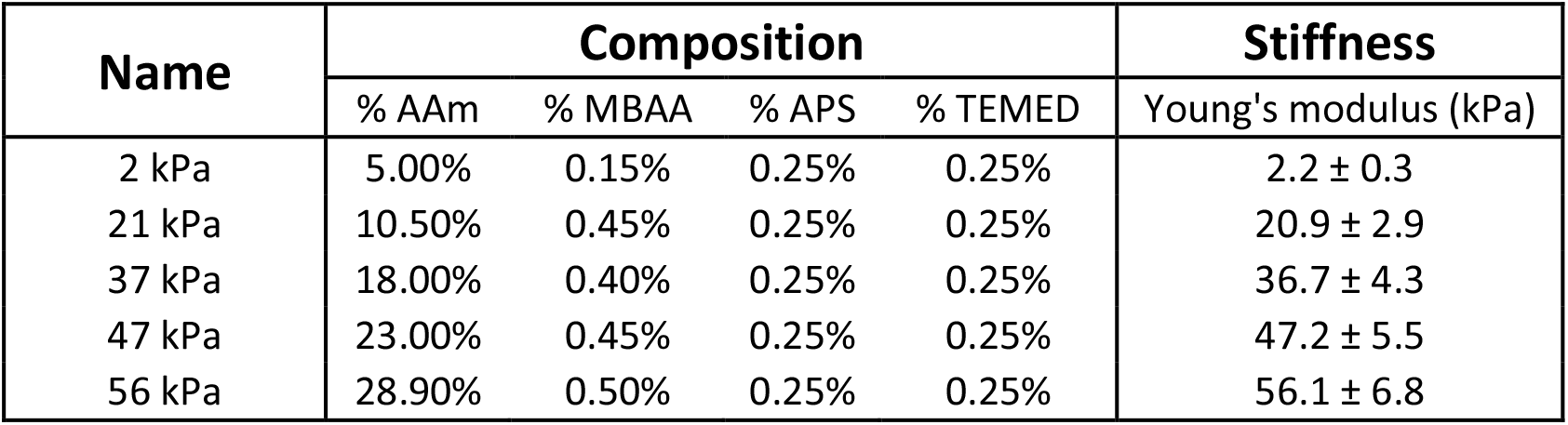
PAAm hydrogel stiffnesses. Gel compositions are given in final concentrations (percentages for AAm, MBAA and APS; V/V ratio for TEMED). Average stiffnesses and errors are measured on a rotational rheometer from minimally three technical replicates. Details of the characterization can be found in Table S1.

#### Hydrogel functionalization

A swollen PAAm hydrogel was restamped into a 75 mm circular gel, washed three times with 10 mL dH_2_O, and incubated under 365 nm UV-light for 15 min with 666 µg sulfosuccinimidyl 6-(4-azido-2-nitrophenylamino)hexanoate (sulfo-SANPAH, Sigma-Aldrich, Merck, 803332) diluted in 10 mL sterile dH_2_O, with a coating density of 0.1 mg/cm^2^. This incubation step was repeated with a new sulfo-SANPAH solution, resulting in a rust-brown colored gel. After three washes with 10 mL sterile dH_2_O to remove unreacted sulfo-SANPAH, the gel was transferred to a new Petri dish, and incubated overnight on a seesaw rocker plate at room temperature with 165 µg collagen type I (Thermo Fisher Scientific, A1048301) diluted in 10 mL sterile 0.5% acetic acid (Acros Organics, 64-19-7, 99.8% solution diluted in dH_2_O) adjusted to pH 3.40 using 1 M NaOH, with a coating density of 25 µg/cm^2^. The acidic pH facilitates a more homogeneous collagen distribution on the gel surface.^38^ Following three washes with 10 mL sterile DPBS to eliminate unreacted collagen, the gel was equilibrated in 10 mL sterile prewarmed sterile cell culture medium in a humidified incubator at 37 °C and 5% CO_2_ for minimally 30 minutes before cell seeding (see Materials and Methods section “cell culture”).

### Cell culture

#### Cell lines

Isogenic KM12C and KM12L4a colorectal cancer cell lines from Fidler’s lab (MD Anderson Cancer Center), human primary colorectal CAF immortalized with hTERT (CT5.3) from De Wever’s lab, and primary green fluorescent Human Umbilical Vein Endothelial Cells (TTFLUOR HUVEC) from Innoprot (Bizkaia, Spain) were used.^22,23,39^ All cell lines, except HUVEC, were grown in standard culture medium, namely Dulbecco’s Modified Eagle Medium (DMEM, Gibco, Thermo Fisher Scientific, 31053028) supplemented with 10% FBS (Sigma-Aldrich, Merck, F7524), 1% GlutaMAX (Gibco, Thermo Fisher Scientific, 35050038) and 0.1% Gentamycin (Carl Roth, 2475.1). HUVEC cells were grown in EBM™-2 Basal Medium (Lonza, CC-3156) supplemented with 2% FBS, hydrocortisone, hFGF-B, VEGF, R3-IGF-1, ascorbic acid, hEGF, GA-1000, and heparin (complete EBM-2 medium) according to the manufacturers’ instructions (EGM™-2 SingleQuots™ Supplements, Lonza, CC-4176). All cells were maintained in a humidified incubator at 37 °C and 5% CO_2_. HUVECs were used up to passage 2.

#### Seeding, growing, and starving cells on PAAm gels

Ten million KM12C or KM12L4a cells were seeded on a 75 mm equilibrated PAAm gel, with the stamp placed around the gel to prevent cells flowing off the gel, resulting in a cell density of approximately 1.5×10^6^ cells/cm^2^. The cell-containing solution was incubated for 20-30 min to enable initial cell attachment and cells were subsequently cultured for 24 h in a humidified incubator at 37 °C and 5% CO_2_. After this initial growth period, cells underwent a 48 h starvation period in serum-free medium to eliminate any interference of serum components in subsequent analyses. To initiate starvation, the gel was washed two times with 10 mL prewarmed commercial DPBS (1x Gibco, Thermo Fisher Scientific, 14190144) and then transferred to a new Petri dish to discard cells that had flowed off the gel and grown on the plastic. The washing steps were repeated, and cells were starved in 13 mL prewarmed starvation medium (standard cell culture medium without 10% FBS) for 48 h in a humidified incubator at 37 °C and 5% CO_2_.

#### Secretome and cell collection

For secretome collection, the medium of starved cells was harvested, centrifuged 5 min at 140 g to remove dead cells, and frozen at -80 °C for further analysis. A portion of this solution (10 mL) was concentrated 10x by freeze-drying and resuspension in 1 mL sterile MilliQ water (MQ) for specific downstream analyses. For cell collection, cells were washed three times with 10 mL prewarmed commercial DPBS and harvested with 10 mL prewarmed 4 mM EDTA (Thermo Fisher Scientific, J15694.AE) diluted in commercial DPBS. To facilitate cell detachment, the solution was flowed over the gel multiple times. Detachment was confirmed under a transmission light microscope before collecting the suspension. In addition, the gel was rinsed once with 3 mL prewarmed commercial DPBS, and this solution was combined with the cell suspension previously collected. Cells were pelleted by centrifugation for 5 min at 140 g, resuspended in 500 µL commercial DPBS for cell counting, and then centrifuged again to obtain a dry cell pellet that was frozen at -80 °C for further analysis.

### Immunofluorescence

To visualize protein expression and subcellular localization, cells grown and starved on top of PAAm gels were immunolabeled following a previously established protocol.^35,40^ Briefly, a 75 mm gel was washed three times with 10 mL prewarmed DPBS and fixed in 10 mL prewarmed paraformaldehyde 4% (PFA, Thermo Fisher Scientific, 28908) diluted in DPBS for 1 h in a humidified incubator at 37 °C and 5% CO_2_. Following fixation, the gel was washed three times with 10 mL prewarmed DPBS and stamped into a circle with 10 mm diameter for immunolabeling. All subsequent washes were performed three times with 200 µL prewarmed DPBS, and all subsequent dilutions were made in 200 µL prewarmed blocking buffer (Table S2) unless stated otherwise. The 10 mm sample was permeabilized in 200 µL prewarmed Triton X-100 0.1% (Sigma-Aldrich, Merck, T8787) for 30 min at 4 °C. The sample was washed, blocked in 200 µL blocking buffer for 2 h in a humidified incubator at 37 °C and 5% CO_2_, and incubated with primary antibody (Table S2) overnight at room temperature. The sample was washed three times for 5 min each, and incubated with secondary antibody (Table S2) diluted in blocking buffer for 2 h at room temperature, protected from light. The sample was washed three times for 5 min each, cell nuclei were stained with 1 µg 4’,6-Diamidino-2-Phenylindole (DAPI, Invitrogen, Thermo Fisher Scientific, D1306) for 15 min in a humidified incubator at 37 °C and 5% CO_2_, washed again three times for 5 min each. To facilitate high-resolution imaging, the sample was placed inverted in a glass-bottom 6-well plate (Cellvis, P06-1.5H-N), with the cells positioned closest to the coverslip, and kept in approximately 1.2 mL DPBS so that the sample was surrounded by liquid but did not float inside the well. Imaging was performed on a Leica TCS SP8 microscope (Leica Microsystems GmbH) with 63x water (NA: 1.2) immersion objective. The sample was excited with a diode laser at 405 nm for DAPI, and at 638 nm for Abberior STAR RED. Emission was captured on highly sensitive hybrid detectors (HyD SMD, Leica Microsystems GmbH), and transmission images were captured simultaneously on a transmitted light detector. Image acquisition was performed by taking z-stacks with 512×512 pixels per field of view, 200 Hz line scan speed and 4-line averaging.

Tubulin signal was quantified using the 3D volume rendering mode of the Imaris Viewer (Oxford instruments). Cell clumps were identified as objects with surface detail set to 0.5, and the manual threshold for low intensity was set to 5, and automatically segmented by the Imaris algorithm. The identified volumes were then filtered based on area and distance to the border, ensuring only clumps fully within the field of view were identified. For each clump, the accumulated DAPI and tubulin fluorescence signal, 3D volume, and sphericity were calculated.

### Protein extraction and quantification

Proteins were extracted from the collected pellets by cell lysis using RIPA buffer (Sigma-Aldrich, Merck, R0278), with 1:100 v/v of proteases and phosphatases inhibitor cocktails (MCE, HY-K0022 and HY-K0010), followed by centrifugation for 10 min at 10,000 g and 4 °C to pellet cell debris and obtain uncontaminated protein extracts. Protein extracts were quantified using the Tryptophan fluorescence method, and protein quality and concentration was confirmed by separating 5 µg of protein extract on a 10% SDS-PAGE gel under reducing conditions alongside a PageRuler Plus Prestained Protein Ladder (Thermo Fisher Scientific, 26619) and Coomassie blue staining.^41^ Secretome solution (900 µL) was directly used as such for TMT labeling.

### TMT labeling

For the TMT analyses, cell extracts and secretome samples obtained after incubation with FBS-free DMEM were used. 10 μg of each protein extract in 100 μL of RIPA or 900 µL of secretome were reduced with 10 mM TCEP for 45 min at 37 °C and 600 rpm and alkylated with 40 mM chloroacetamide for 30 min at room temperature, 600 rpm, protected from light. Three replicates per condition were analyzed (R1-3).

Protein extracts were incubated with 100 µL of SeraMag magnetic beads mix (50% hydrophilic beads - 50% hydrophobic beads, GE Healthcare, Chicago, Illinois, USA) and 200 μL of acetonitrile (ACN) for 35 min at room temperature and 600 rpm for protein binding to the beads. Supernatants were then discarded, and magnetic beads were washed twice with 70% ethanol and once with ACN. Finally, supernatants were discarded, and proteins were digested overnight at 37 °C and 600 rpm with 1:20 porcine trypsin (Thermo Fisher Scientific) - 0.5 μg - in 100 μL of 200 mM HEPES, pH 8.0. For the analysis of the secretome, samples were incubated with 100 µL of SeraMag magnetic beads mix and then divided into two tubes and incubated with 500 μL ACN for protein binding to the beads. Beads were processed as previously described but pooling together the split beads after the first wash with 70% ethanol. The following day, samples were sonicated twice, supernatants were collected and labeled separately with eighteen different Tandem Mass Tags (TMT) reagents (Thermo Fisher Scientific). TMT reagents were previously resuspended 21 µL ACN, and samples were labeled in two incubation steps of 30 min at room temperature and 600 rpm and with 10 μL of reagent per incubation. Samples were then incubated with 10 μL of 1 M glycine, pH 2.7, for 30 min at room temperature and 600 rpm. The contents of the 18 tubes were pooled together and dried under vacuum prior to peptide separation using the High-pH Reversed-Phase Peptide Fractionation Kit (Thermo Fisher Scientific). Briefly, desiccated peptides were reconstituted in 300 μL of 0.1% TFA in MQ, and columns were equilibrated twice with 300 μL of ACN and twice with 300 μL of 0.1% TFA in MQ. The peptide solutions were loaded into the columns, washed twice with 300 μL of 0.1% TFA in MQ, and separated in 12 fractions of 300 µL each in 0.1% triethylamine and 2.5-100% ACN. Fractions were then dried under vacuum, and stored at -80 °C until analysis in twelve LC-MS/MS runs. Samples were reconstituted in 10 μL of 0.1% FA prior to their injection onto the LC-MS/MS mass spectrometer.

### LC-MS/MS analysis

TMT experiments were analyzed on an Orbitrap Exploris 480 mass spectrometer (Thermo Fisher Scientific) equipped with the FAIMS Pro Duo interface (Thermo Fisher Scientific). Peptide separation was performed on the Vanquish Neo UHPLC System (Thermo Fisher Scientific). For each analysis, samples were loaded into a precolumn PepMap 100 C18 3 µm, 75 µm × 2 cm Nanoviper Trap 1200BA (Thermo Fisher Scientific) and eluted in an Easy-Spray PepMap RSLC C18 2 µm, 75 µm × 50 cm (Thermo Fisher Scientific) heated at 50 °C. The mobile phase flow rate was 300 nL/min, and 0.1% FA in MQ and 0.1% FA in 80% ACN were used as buffers A and B, respectively. The 2 h gradient was: 0-2% buffer B for 4 min, 2% buffer B for 2 min, 2-42% buffer B for 100 min, 42-72% buffer B for 14 min, 72-95% buffer B for 5 min, and 95% buffer B for 10 min. Samples were resuspended in 10 µL of buffer A, and 2-4 µL (800 ng) of each sample were injected per run. For ionization, 1900 V of liquid junction voltage and 280 °C capillary temperature were used. The full scan method employed a m/z 350-1400 mass selection, an Orbitrap resolution of 60,000 (at m/z 200), an automatic gain control (AGC) value of 300%, and a maximum injection time (IT) of 25 ms. After the survey scan, the 12 most intense precursor ions were selected for MS/MS fragmentation. Fragmentation was performed with a normalized collision energy of 32, and MS/MS scans were acquired with a 100 m/z first mass, an AGC target of 100%, a resolution of 15,000 (at m/z 200), an intensity threshold of 2×10^4^, an isolation window of 0.7 m/z units, a maximum IT of 22 ms, and the TurboTMT enabled. Charge state screening was enabled to reject unassigned, singly charged, and greater than or equal to seven protonated ions. A dynamic exclusion time of 30 s was used to discriminate against previously selected ions. A gas flow of 4 L/min and CVs = -45 V and -60 V were used for FAIMS.

MS data were analyzed with MaxQuant (version 2.4.2, Max Planck Institute of Biochemistry, Planegg, Germany) using standardized workflows. Mass spectra ^*^.raw files were searched against the Uniprot UP000005640_9606.fasta *Homo sapiens* (human) 2022 database (20,577 protein entries) using reporter ion MS2 type for TMTs. Precursor and reporter mass tolerances were set to 4.5 ppm and 0.003 Da, respectively. Trypsin was set as the digestion enzyme, allowing a maximum of 2 missed cleavages per peptide. Carbamidomethylation of cysteines was set as a fixed modification, and methionine oxidation, N-terminal acetylation, and Ser, Thr, and Tyr phosphorylation were set as variable modifications. Unique and razor peptides were considered for quantification. Minimal peptide length and maximal peptide mass were fixed to 7 amino acids and 4600 Da, respectively. Identified peptides were filtered by their precursor intensity fraction (PIF) with a false discovery rate (FDR) threshold of 0.01. Proteins identified with at least one unique peptide and an ion score above 99% were considered for evaluation, whereas proteins identified as potential contaminants were excluded from the analysis. The protein sequence coverage was estimated for specific proteins by the percentage of matching amino acids from the identified peptides having confidence greater than or equal to 95% divided by the total number of amino acids in the sequence. In addition, reporter ion intensities were bias-corrected for the overlapping isotope contributions from the TMT reagents according to the manufacturer’s certificate.

Data normalization was performed to equalize the differences in the total sum of signals for each TMT channel, as the same amount of protein was labeled in each TMT sample. Sample loading (SL) normalization was performed with R Studio (version 4.1.1, Posit PBC, Boston, Massachusetts, USA) according to the established protocol (https://github.com/pwilmart, accessed on November 2^nd^ 2022), using the “tidyverse”, “psych”, “gridExtra”, “scales”, and “ggplot2” packages.

For statistical analysis, an empirical Bayes-moderated t-statistics analysis was performed with R Studio (version 4.1.1) using the packages “limma”, “dplyr”, “tidyverse”, “ggplot2”, and “rstatix”, according to previously described procedures.^42–45^ Prior to statistical analysis, reverse and contaminant proteins were removed, and data filtering (proteins identified in at least 60% of samples were considered for the analysis) and missing value imputation by random draws from a Gaussian using the “imputeLCMD” R package were performed. Proteins identified with one or more unique peptides, an expression ratio ≥ 1.5 (upregulated) or ≤ 0.67 (downregulated), and a false discovery rate (FDR) ≤ 0.05 were selected as statistically significant dysregulated proteins. Expression ratio cutoffs were selected according to previous reports.^32,46–48^ Venn diagrams were obtained using the Jvenn website (http://bioinfo.genotoul.fr/jvenn, accessed on 21st May 2023). Heatmap and GeneOntology and Reactome analyses of significantly dysregulated proteins was performed with R Studio (version 4.1.1) using the “pheatmap”, “stringdb”, “fgsea”, “ReactomePA”, and “biomaRt” packages and the Reactome Pathway Database (https://reactome.org/PathwayBrowser/#/, accessed on May 21^st^, 2023).

To determine secretion pathways of dysregulated 2 kPa *versus* 47 kPa proteins, SecretomeP 2.0 and SignalP 6.0 were employed, similar to our previous analysis.^49^ Briefly, secretion of proteins was evaluated using SecretomeP 2.0 with 0.6 as cutoff value for proteins of eukaryotic origin, as recommended by the application.^50^ Secreted proteins were further analyzed with SignalP 6.0, where a value > 0.45 indicated the presence of a signal peptide, reflecting classical secretion. Finally, the ExoCarta database was used to identify non-classically secreted proteins in previously identified human exosomes.^51–54^

### Dot blot

For validation of the dysregulation of selected proteins, we performed a dot blot with the 6 replicates of KM12L4a cells (KM12L4a R1-6) and the 2 replicates of KM12C cells (KM12C R1-2). For dot blot, 5 µL or 25 µL non-lyophilized secretome diluted in DPBS to a total volume of 100 µL, was blotted onto a nitrocellulose membrane using the Bio-Dot 96-Well Microfiltration Apparatus (Bio-Rad, 72926-1). Protein transfer was verified by Red Ponceau staining (Sigma-Aldrich, Merck, P3504). Membranes were blocked in blocking buffer (Table S2) on a seesaw rocker plate for 1 h at room temperature, and incubated with primary antibody diluted in blocking buffer (Table S2) on a seesaw rocker plate overnight at 4 °C. Then, membranes were washed three times with 0.1% Tween 20 diluted in DPBS for 10 min each, and incubated with the appropriate HRP-conjugated secondary antibody (Table S2) diluted in blocking buffer on a seesaw rocker plate for 1 h at room temperature. Membranes were washed three times with 0.1% Tween 20 diluted in DPBS for 10 min each, and the signal was developed using the ECL Pico PLUS Chemiluminescent Substrate (Thermo Fisher Scientific, 34580) or, alternatively, the SuperSignal West Femto Maximum Sensitivity Substrate (Thermo Fisher Scientific, 34096). Images were recorded on an Amersham ImageQuant 800 (Cytiva). For image quantification, the integrated density inside an equally sized circular region was measured for the different dots in Fiji (ImageJ), simultaneously analyzing 2 and 47 kPa dots from the same replicate.^55^ For each replicate, the image with optimal exposure conditions was analyzed and data was plotted as the ratio of 47 *versus* 2 kPa.

### Cytokine array

Secretome from 2 and 47 kPa non-lyophilized samples were directly analyzed using the Human Cytokine Antibody Array C5 kit (RayBiotech, AAH-CYT-5-8), according to the manufacturers’ instructions. Three replicates were analyzed per condition (KM12L4a R1-3). Briefly, 0.9 mL secretome was incubated overnight at 4 °C on an antibody microarray previously blocked with the supplied blocking buffer. After the washing steps, the biotinylated antibody cocktail was incubated overnight at 4 °C. After new washing steps, HRP-streptavidin was incubated for 2 h at room temperature. After final washing steps, the chemiluminescence detection was developed and recorded immediately afterwards on an Amersham ImageQuant 800 (Cytiva). Signal quantification was performed with the gel analysis tool in Fiji (ImageJ), simultaneously analyzing corresponding rows of aligned 2 and 47 kPa membranes from the same replicate.^55,56^ For each spot, the image with optimal exposure conditions was analyzed and data was plotted as the ratio of 47 *versus* 2 kPa.

### ELISA

For validation of the results obtained by antibody microarrays, secretome from 2 and 47 kPa non-lyophilized samples were surveyed by ELISA, according to the manufacturer’s instructions, to determine the protein levels of VEGF, IL-8, CXCL1, and osteoprotegerin (OPG) in 6 replicates of KM12L4a cells (KM12L4a R1-6) and 2 replicates of KM12C cells (KM12C R1-2). Human IL-8 sandwich ELISA kit (Proteintech, KE00006) and human CXCL1 sandwich ELISA kit (Proteintech, KE00133) were used with 25 µL secretome samples diluted 1:2 in PT 4-eg sample diluent and PT 1-ef sample diluent (provided in the kit), respectively. Additionally, 5 µL and 10 µL secretome samples 1/10 and 1/5 diluted in sample diluent (provided in the kit) were analyzed using the human VEGF sandwich ELISA kit (Proteintech, KE00216) and the human OPG ELISA kit (Elabscience, E-EL-H1341), respectively. Color was developed according to the manufacturer instructions and signal recorded onto a Spark multimode microplate reader (Tecan Trading AG, Männedorf, Switzerland).

### Functional assays

#### Wound healing

KM12C cells were seeded into wound healing inserts (Ibidi, #80209) attached to a plastic-bottom 24-well plate (Sarstedt, 83.3922). Cells were seeded at a density of 1.5×10^5^ cells per insert of the well (3×10^5^ in total per well). The next day, the inserts were removed to uncover a cell-free area of 500 ± 100 µm. 500 µL of fresh cell growth medium was then added to the wells, along with 5 µL of the corresponding lyophilized secretomes (KM12L4a R1 and R4) or control (lyophilized starvation medium). Time-lapse imaging of the cells was recorded using a Leica THUNDER imager (Leica Microsystems GmbH). Images (1024×1024 pixels, 1327×1327 µm) were acquired every hour under constant incubation conditions (37 °C and 5% CO_2_), using a 10x air objective (NA: 0.32). Time-lapse images were analyzed using the MRI’s Wound Healing tool for Image J software (NIH) (https://github.com/MontpellierRessourcesImagerie/imagej_macros_and_scripts/wiki/Wound-Healing-Tool). We visually inspected all calculated cell-free areas to ensure accurate determination. The wound closure speed was calculated by plotting the cell-free area over time and determining the slope on the linear segment of the plot.

#### Angiogenesis

A 15-wells µ-Slide angiogenesis chamber (Ibidi, 81506) was coated with 10 µL Matrigel (Merck, CLS356234) for 1 h at 37 °C and 5% CO_2_. TTFLUOR HUVECs were harvested with trypsin and centrifuged for 5 min at 140 g. 2×10^4^ cells were resuspended in 25 µL complete EBM-2 medium and 25 µL of lyophilized secretome (without FBS) diluted in sterile DPBS (1.5 µL lyophilized secretome in 23.5 µL DPBS) were added per condition. The positive control only contained complete EBM-2 medium. The mixture was added onto the Matrigel-coated coverslip. Tube formation was monitored immediately after seeding though time-lapse imaging using a Leica THUNDER imager (Leica Microsystems GmbH). Images (1024×1024 pixels, approximately 1325×1325 µm) were acquired in the GFP channel every 5 minutes during 6 h, for 9 different fields of view per condition, under constant incubation conditions (37 °C and 5% CO_2_), using a 10x air objective (NA: 0.32). Because a dilution with the different lyophilized secretome samples also results in a dilution of the angiogenic factors of the complete EBM-2 medium, this might reduce the tube formation capacity of HUVEC cells. Tube formation was analyzed in Fiji on ROIs of equal size, by enhancing the contrast and subtracting the background, applying a median filter with a 3-pixel radius, followed by an automatic threshold and two iterations of erosion, and segmenting the cells based on a watershed algorithm.^55^ The area of individual features was quantified and normalized by division of the mean area at timepoint 0. The increment change of this normalized area over time was determined based on a linear fit in Excel, and normalized to the median of the control condition.

#### Gelatin degradation

Coverslips coated with fluorescently labeled gelatin were prepared based on a protocol kindly provided by Prof. E. Planus.^57,58^ Briefly, 12 mm round glass coverslips (VWR, 631-1577) were sequentially washed in absolute ethanol and dH_2_O, then dried under a biological safety cabinet, sterilized under 365 nm UV-light for 15 min, and placed in a glass-bottom 24-well plate (Cellvis, P24-1.5H-N) with one coverslip per well. These coverslips were coated with 25 µg poly L-lysine (Sigma-Aldrich, Merck, P8920) diluted in 500 µL sterile dH_2_O, incubated for 20 min at room temperature, and washed twice with sterile dH_2_O and once with sterile DPBS. Fixation was performed with 500 µL 0.5% glutaraldehyde (Alfa Aesar, A17876) diluted in ice cold sterile DPBS, for 15 min on ice, followed by three washes with ice cold DPBS. Coverslips were then incubated with a mixture of fluorescently labeled gelatin. The gelatin mixture was prepared by dissolving aliquots of non-fluorescent gelatin (Sigma-Aldrich, Merck, G1890, dissolved in DPBS to 2 mg/mL) and FITC-conjugated gelatin (AnaSpec, AS-85145, dissolved in dH_2_O to 8 mg/mL) separately at 45 °C for 15 min. Both gelatins were then mixed in sterile DPBS to a final concentration of 0.2 mg/mL for each component. 500 µL of this mixture was incubated on a coverslip for maximally 10 min at room temperature. After incubation, coverslips were washed three times with 2 mL sterile DPBS, and sterilized under 365 nm UV-light for 5 min. For cell seeding, CAF CT5.3 cells were harvested and labeled with cell tracker red CMTPX (Thermo Fisher Scientific, C34552). Briefly, 1×10^5^ cells were centrifuged 5 min at 140 g and resuspended in 0.15 µL cell tracker red CMTPX (1:500 dilution in starvation medium, standard cell culture medium without FBS). Cell tracker was incubated for 30 min in a humidified incubator at 37 °C and 5% CO_2_, with resuspension every 10 min to avoid cell clumping. Unbound cell tracker was removed by centrifugation, 1×10^4^ cells were resuspended in 500 µL standard medium with 5 µL lyophilized secretome (KM12L4a R1 and R4-6) or control (lyophilized starvation medium), and seeded on a gelatin-coated coverslip. After incubation for 48 h in a humidified incubator at 37 °C and 5% CO_2_, gelatin degradation was evaluated by imaging on a Leica TCS SP8 microscope (Leica Microsystems GmbH) with a 20x air objective (NA: 0.75). Images were taken as z-stacks of 512×512 pixels, with sequential acquisition in the FITC and cell tracker red channel with excitation at 488 nm and 552 nm, respectively. Images were quantified in Fiji by making a maximum z-projection of the cell tracker channel, performing an intensity-based segmentation based on visual observations and using a black background, and using the dilate (once) and erode (twice) commands to retrieve an uninterrupted outline of the cell.^55^ Then, the average intensity value of the gelatin channel was measured on the z-projection of the gelatin-FITC channel, both inside and outside the cell mask. The ratio for inside *versus* outside was calculated, meaning that values below 1 reflect gelatin degradation inside the cell.

### Statistical analyses

Statistical analyses were performed in GraphPad Prism 10 (GraphPad), except when stated otherwise. For statistical analyses of dot blots, cytokine arrays, and ELISAs, a one-sided t-test with unequal variances and a hypothetical mean of 1, was performed in Microsoft Excel. For 3D cell morphology and cytoskeleton analyses, unpaired t-tests were performed to assess significant differences, either using a parametric test with Welch’s correction for normally distributed data or using a nonparametric Mann-Whitney test. For wound healing and angiogenesis analyses, one-way ANOVAs were performed to assess significant differences. For gelatin degradation, statistical significances were determined using a nonparametric Kruskal-Wallis test, comparing mean ranks of the data groups in an uncorrected Dunn’s test. All significances were evaluated as follows: non-significant difference for p > 0.05 (-), significant difference for p ≤ 0.05 (^*^), significant difference for p ≤ 0.01 (^**^), significant difference for p ≤ 0.001 (^***^), significant difference for p ≤ 0.0001 (^****^).

## Results

### 1. Replicating CRC stiffnesses with PAAm hydrogels

To mimic the changes in ECM stiffness observed in CRC progression, we developed a library of synthetic polyacrylamide (PAAm) hydrogels. These hydrogels, produced as large, free-standing gels, are easily customizable in both shape and size, rendering them ideally suited for different applications, namely proteomic analysis and multiplexed immunolabeling.^34,35^ The stiffness of the synthesized PAAm gels was tuned by varying the acrylamide (AAm) and bisacrylamide (MBAA) content, and measured using a rotational rheometer (Table 1, and Materials and Methods and Table S1 for details). The softest gel (2 kPa) mimics the stiffness of healthy colorectal tissue, whereas the stiffer gels (21-56 kPa) simulate the stiffness of advanced CRC (stages T3 and T4).^7^

Culturing metastatic KM12L4a CRC cells on collagen functionalized PAAm gels and collagen-coated glass (control) revealed substantial differences in cell morphology depending on the substrate stiffness. Cell aggregates formed under all conditions, but exhibited more protrusions and interconnections on stiffer substrates and on the glass control, while remaining more rounded on softer 2 and 21 kPa substrates (Fig. 1A). To quantify these differences, we focused on two stiffness values (2 and 47 kPa) with clear differences in cell spreading. To assess if cytoskeletal changes were driving the observed morphological differences, we visualized 3D cell structure through immunolabeling of tubulin, a major component of the cytoskeleton (Fig. 1B). Consistent with our initial observations, we found more and smaller aggregates on the softer substrates (Fig. 1Ci-iii), with cells forming tightly packed, spherical aggregates (Fig. 1Civ-v). On stiffer substrates, particularly at the aggregate edges, cells spread out and exhibited cytoskeletal protrusions, even though the tubulin expression level remain similar to that on the softer substrate (Fig. 1Cvi). These results indicate that substrate stiffness influences cell morphology, underscoring that KM12L4a cells are mechanosensitive and therefore suitable for studying the impact of altered ECM stiffness.

**Figure 1:**
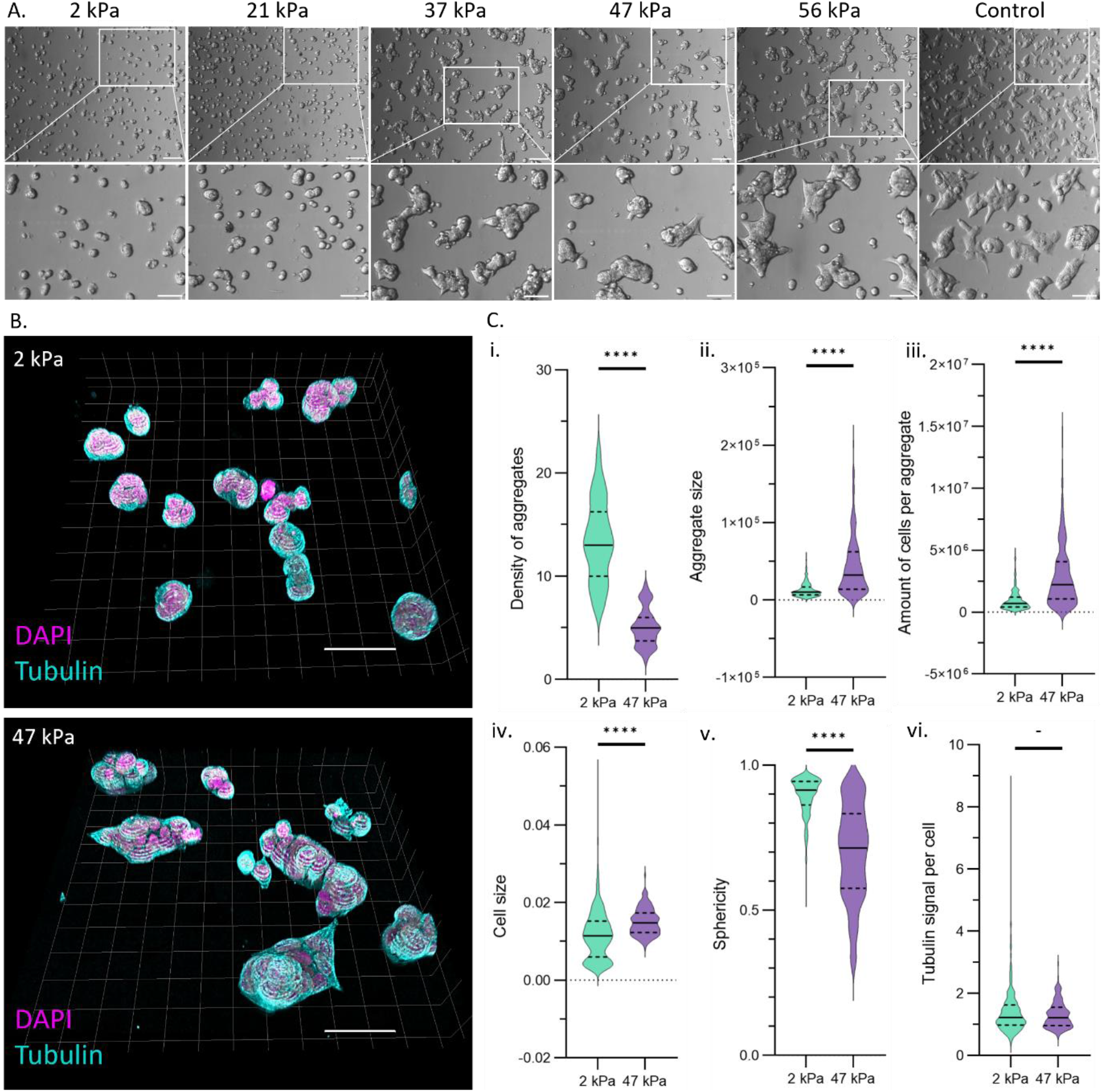
Growth of metastatic KM12L4a cells on top of collagen functionalized PAAm hydrogels, and collagen-coated glass (control). Images are representative of cells before proteomic analysis: after 24 h cell growth, 48 h starvation and three washes with DPBS. **A:** Representative transmission images. Scale bars: 200 µm and 100 µm, for full size and magnifications, respectively. **B:** Representative 3D view of tubulin (cyan) and DAPI (magenta) in KM12L4a cells grown on 2 and 47 kPa hydrogels. Grid represents 3D-view, with XY scale bars: 50 µm. **C:** 3D morphological quantification for different parameters. Data is represented in a violin plot where solid lines show medians, while dashed lines show quartiles. Data was obtained from n = 30 samples, from three biological replicates. Significance levels: (-) p > 0.05, non-significant; (^****^) p ≤ 0.0001. Images in panels A and B are from different samples.

### 2. Exploring stiffness-induced changes on protein expression

#### 2.1 In-depth quantitative TMT-based LC-MS/MS analysis

To investigate how substrate stiffness affects protein expression, we analyzed both cellular and secreted proteins by LC-MS/MS. KM12L4a cells were cultured for 72 hours on collagen functionalized PAAm gels and on collagen-coated glass (control). During the final 48 hours, the standard cell culture medium was replaced by serum-free medium to avoid FBS interference in the proteomic analysis. The experiments, performed in triplicate (R1-3), included PAAm hydrogels with five different stiffnesses and the glass control, resulting in 18 secretome and 18 cell samples. Cell proteins were extracted by cell lysis with RIPA buffer and sample quality was verified by Coomassie blue staining prior to TMT-labeling (Fig. S1). Secretome samples were reduced, alkylated, and TMT-labeled directly.

After cell lysis, each cell extract and secretome sample was separately digested using trypsin, labeled with 18 different TMT reagents, and analyzed by LC-MS/MS. After normalizing the data (Fig. S2), 6484 proteins from the cell extracts and 3333 proteins from secretome samples were identified and quantified with at least one unique peptide and an ion score above 99% (Tables S3-S4). Principal component analysis (PCA) was then performed to investigate the variability among biological replicates and to compare the protein expression profile of cells cultured on the different stiffnesses. No clear separation was observed between the cell extracts of samples grown on different substrates (Fig. S4A). In contrast, the TMT proteomic analysis of the secretome showed a clear separation with increasing substrate stiffness (Fig. 2A). The secreted protein profiles for 2 kPa and glass conditions were well-separated from each other and from the other stiffnesses. The 21 – 56 kPa conditions clustered according to increasing stiffness, except for 37 and 47 kPa, which grouped closely together.

**Figure 2:**
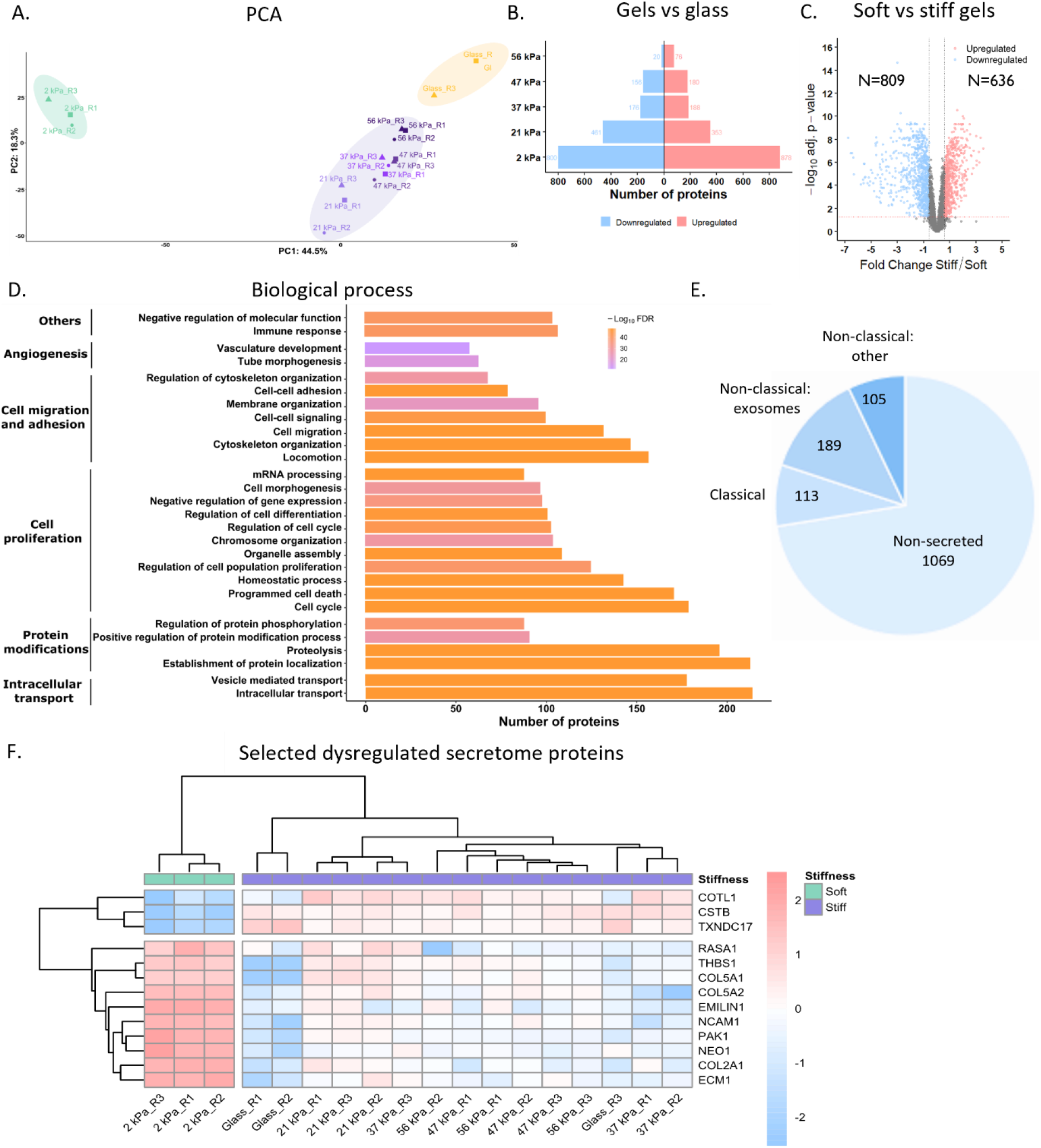
Proteomic analyses of secretome samples. **A-C:** Differential expression analyses. **A:** Principal component analyses (PCA) representing sample clustering according to the stiffness. **B:** Bar plots with the number of upregulated (red) and downregulated (blue) proteins in the different stiffness conditions, compared to glass. Volcano plots with the fold changes and p-values are shown in Fig. S3A. **C:** Volcano plots of proteins identified and quantified as altered in stiff (21, 37, 47, and 56 kPa) *versus* soft (2 kPa) conditions. The x-axis represents the Log_2_ expression ratio (fold change) of stiff *versus* soft relative protein expression ratios, the y-axis depicts the FDR value (adj. p-value) based on -Log10. Colored dots represent differentially expressed proteins upregulated (red) and downregulated (blue) in stiff conditions with FDR ≤ 0.05 (represented by red dashed horizontal line) and 1.5-fold expression ratio (represented by two black dashed vertical lines). **D-F:** Bioinformatics analysis of dysregulated proteins in stiff (21, 37, 47, and 56 kPa) *versus* soft (2 kPa) conditions. **D:** Bar plots represent the most significantly enriched biological processes in which dysregulated proteins are involved according to Gene Ontology (GO) analysis. Cellular components and Reactome analysis are shown in Fig. S3B-C. **E**: Location and secretion pathway analysis of dysregulated secretome proteins according to SecretomeP, SignalP and ExoCarta databases. **F**: Heatmap of the TMT relative protein expression of the 12 dysregulated secretome proteins selected for validation normalized to the mean row intensity. While 13 dysregulated proteins are shown in the heatmap, COL5A1 and COL5A2 were taken together in our validation assays, as the antibody used binds to all three COL5 isoforms.

#### 2.2 Dysregulation analysis reveals differences between soft and stiff substrates

For further analysis, proteins were considered as dysregulated in cell extracts or secretomes if their expression ratio was ≥ 1.5 (upregulated) or ≤ 0.67 (downregulated), and false discovery rate (FDR) was ≤ 0.05. A significant dysregulation was observed in the secretome of KM12L4a cells cultured on different stiffnesses compared to glass (Fig. 2B, S3A and Table S3), whereas cell extracts showed almost no dysregulation (Fig. S4B and D and Table S4). Interestingly, despite the low number of dysregulated proteins in cell extracts, both the cell extracts and the secretome showed that a higher number of proteins were dysregulated on the softer hydrogel compared to glass, i.e. the stiffest condition. These results suggest that culturing cells on substrates with different stiffnesses, mimicking CRC development and progression, causes the cells to release different factors.

We proceeded by comparing the 2 kPa condition (mimicking healthy colorectal tissue, soft condition) to the four PAAm hydrogels with higher stiffnesses (21 – 56 kPa, advanced CRC stages T3 and T4, stiff conditions). Cell extracts showed minimal dysregulation, with only 3 upregulated and 3 downregulated proteins (Fig. S4C). In contrast, the secretome exhibited extensive dysregulation between soft and stiff conditions, with 636 upregulated and 809 downregulated proteins (Fig. 2C). Therefore, our subsequent analysis primarily focused on the dysregulated proteins within the secretome.

The Gene Ontology (GO) database was used to identify the cellular components and biological processes related to the 1445 dysregulated secretome proteins (Fig. 2D and S3B). These proteins were associated with intracellular transport mechanisms, protein alterations, cell proliferation, cell migration and adhesion, and angiogenesis (Fig. 2D). Cellular component analysis revealed that dysregulated proteins were present in various cellular locations, with a substantial number associated with vesicles, the extracellular matrix and cell attachment (Fig. S3B). Analysis of the dysregulated proteins using the Reactome database identified significant alterations in pathways involved in neutrophil degranulation (83 proteins), N-linked glycosylation (57 proteins), translation (56 proteins), mRNA splicing and rRNA processing (45 and 44 proteins, respectively), signaling by ROBO receptors (45 proteins), and programmed cell death (43 proteins) (Fig. S3C).

### 3. Validating and extending secretome dysregulation analysis

Given the substantial changes observed in the secretome in response to substrate stiffness, several orthogonal techniques were used to validate protein dysregulation between the secretomes of cells grown on 2 and 47 kPa gels (mimicking healthy tissue and advanced CRC stages, respectively).

#### 3.1 In silico analysis elucidates protein secretion pathways

To verify that the dysregulated secretome proteins were not experimental artifacts, we used SecretomeP for an *in silico* analysis to determine the cellular pathways involved in their secretion. This tool estimates the probability that a protein is secreted based on its post-translational modifications and cellular localization. Out of 1476 proteins dysregulated, 407 were predicted to be secreted, but only 113 contained a signal peptide for the classical secretory route, indicating that most of the dysregulated proteins were released into the extracellular environment via alternative pathways (Fig. 2E and Table S5). Analysis with ExoCarta, a database of proteins identified in exosomes from multiple organisms, revealed that 189 proteins can be secreted through exosomes, 68 of which have been associated with colorectal cancer cells. Experimentally, our analysis is supported by previous studies, which identified some of these proteins on the cell surface or in the secretome of KM12 cells, suggesting that they can indeed be released by cleavage or secretion.^49,59^

#### 3.2 Dot blot analyses complement and validate proteomics results

To support our proteomic results, we further investigated the secretome dysregulation by dot blot, cytokine arrays and ELISA tests. Among the most dysregulated secretome proteins between different hydrogel conditions (soft versus stiff) and glass, we selected 12 proteins associated with cell adhesion, cell cycle, or CRC: three upregulated (COTL1, CSTB, and TXNDC17), and nine downregulated (RASA1, THBS1, COL5A1/COL5A2, EMILIN1, NCAM1, PAK1, NEO1, COL2A1, and ECM1).^29,31,32,49,60^ Unsupervised cluster analysis showed significant dysregulation of all 12 secretome proteins in 2 kPa gels compared to the other stiffnesses (Fig. 2F). Remarkably, in the hierarchical cluster analysis of the 6 dysregulated proteins of the cell extracts, the 2 kPa condition was separated from other stiffnesses, but replicates of different stiff conditions clustered randomly (Fig. S4E).

Dysregulation of the 12 selected secretome proteins was analyzed using dot blot analysis (Fig. 3A and S5). The initial biological replicates (KM12L4a R1-3) were analyzed to validate the TMT data, and three additional replicates (KM12L4a R4-6) were included to confirm reproducibility of our results. Protein dysregulation between stiffer (47 kPa) and softer (2 kPa) conditions matched LC-MS/MS results for 10 of the selected proteins (COTL1, CSTB, RASA1, PAK1, EMILIN1, THBS1, COL2A1, COL5, ECM1, and NCAM), although some replicates showed non-significant trends and large error intervals. The other two selected proteins, TXNDC17 and NEO1, showed conflicting dysregulation trends between the original and additional replicates – TXNDC17 upregulation was observed only in the original replicates (R1-3), while NEO1 downregulation was seen in the additional replicates (R4-6) – making it challenging to draw conclusions for these two proteins.

**Figure 3:**
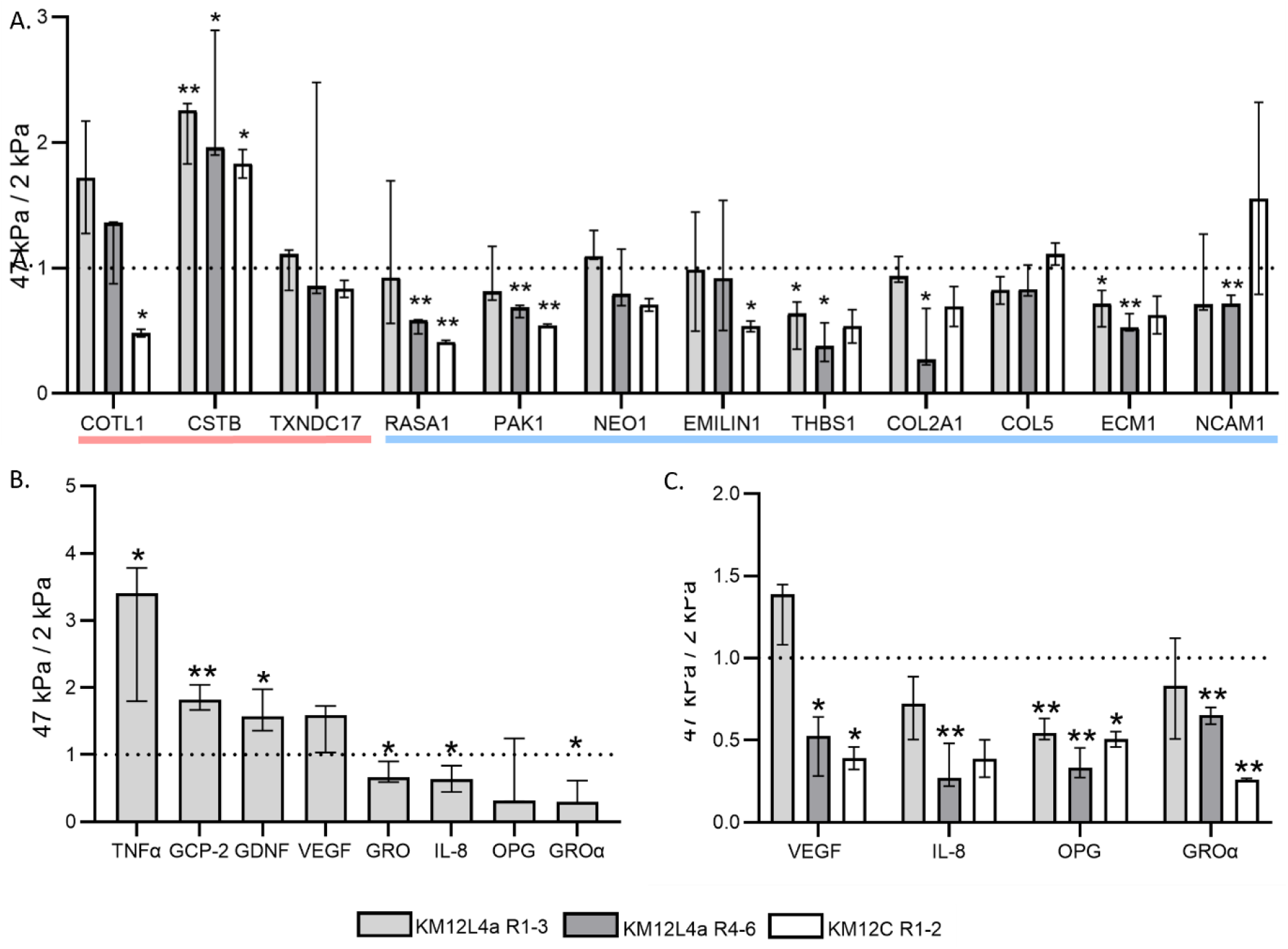
Validating and extending secretome dysregulation analysis for stiffer (47 kPa) versus softer (2 kPa) conditions. **A:** Dot blot analysis of selected proteins found upregulated (red line) and downregulated (blue line) in MS/LC analysis. Full dataset is shown in Fig. S5. **B-C:** Cytokine analyses. **B:** Cytokine array analysis showing upregulated and downregulated cytokines. Dataset is shown in Fig. S6. **C:** ELISA tests for 4 selected cytokines. Data was averaged over two technical replicates. **A-C:** Bars represent median with 95% confidence interval. “R” denotes “replicate”, showing original KM12L4a replicates (R1-3), additional replicates (R4-6) and poorly metastatic KM12C replicates (R1-2). Significance levels: (^*^) p ≤ 0.05; (^**^) p ≤ 0.01.

Two additional replicates from the poorly metastatic KM12C cells (KM12C R1-2) were tested to explore the interplay between metastatic potential and substrate stiffness (Fig. 3A and S5). All selected proteins followed the same trend as the original KM12L4a replicates, except COTL1, COL5, and NCAM. Notably, unlike the ambiguous results of the KM12L4a replicates, the poorly metastatic replicates (KM12C R1-2) confirmed the downregulation of NEO1, as expected from the LC-MS/MS findings. Overall, the dysregulation observed in LC-MS/MS was validated for most of the selected proteins, with additional replicates from the highly metastatic KM12L4a and poorly metastatic KM12C generally showing consistent results.

#### 3.3 Extending proteomics analysis by investigating cytokines

Cytokines, essential for cellular communication, are difficult to detect via LC-MS/MS due to their small size and low abundance in the secretome. To overcome this limitation, we extended our TMT-based proteomic findings to evaluate the expression levels of 80 human cytokines using a cytokine antibody microarray. We compared the softer (2 kPa) and stiffer (47 kPa) conditions in the three original KM12L4a replicates (KM12L4a R1-3). In the secretomes obtained from stiffer conditions, three cytokines were significantly upregulated (TNFα, GCP-2, and GDNF) and three were significantly downregulated (GRO, IL-8, and GROα, Fig. 3B and S6B-C). VEGF and OPG appeared to be up- and downregulated, respectively, but these changes were not statistically significant.

The results from the cytokine array were validated with ELISA tests on selected cytokines: one upregulated (VEGF) and three downregulated (GROα, OPG, and IL-8). These tests were performed in duplicate on the six KM12L4a replicates (KM12L4a R1-6) and on two KM12C replicates (KM12C R1-2) (Fig. 3C). While some comparisons between stiffer and softer conditions were not significant due to the sample size, GROα and IL-8 were consistently downregulated for all replicates, in line with the cytokine array. ELISA tests for VEGF and OPG, which showed non-significant differences in the cytokine arrays, confirmed non-significant changes for VEGF (for KM12L4a R1-3), but revealed significant changes in OPG across the KM12L4a replicates (KM12L4a R1-6). Notably, IL-8, OPG, and GROα trends were consistent across highly metastatic (KM12L4a R1-6) and poorly metastatic (KM12C R1-2) cells, but VEGF showed the opposite trend between KM12L4a R1-3 (upregulated) and the other replicates (KM12L4a R4-6 and KM12C R1-2, downregulated).

Despite their low abundance, nine of the tested cytokines were identified in the LC-MS/MS analysis (Fig. S6B-C and E). Four cytokines (VEGF, IGFBP-2, OPG, TIMP-2) showed similar trends in both methods, although the differences were not always significant, and the results for four cytokines (angiogenin, IGFBP-3, IGFBP-4, TIMP-1) were inconclusive due to large differences between replicates. TGF-β1 showed opposite results: upregulated in cytokine arrays, downregulated in LC-MS/MS.

### 4. Analyzing functional effects of altered secretome

Our analysis identified a dysregulation in several secreted factors that can influence or regulate key cellular processes, including inflammation (e.g. GRO-family, interleukin 8), cell growth and apoptosis (e.g. TGFβ-family), cell migration and metastasis (e.g. GCP2, GDNF), angiogenesis (e.g. VEGF, ANG, THBS1), and matrix remodeling (MMPs). In the final part of this study, we used cell-based assays to evaluate how the different secretomes impact cellular behaviour *in vitro*.

#### 4.1 Effects on cell migration

Understanding the mechanisms driving cancer cell migration is crucial, as it plays a pivotal role in tumor invasion and metastasis. Given the observed dysregulation of factors involved in cell migration and adhesion, we investigated how the secretome influences the migratory behavior of cancer cells. Wound healing assays revealed that the migration of poorly metastatic KM12C cells is affected by the KM12L4a secretome (Fig. 4A). Compared to the control (lyophilized starvation medium), the secretome from KM12L4a cells cultured on softer substrates (2 kPa) increased the migration speed of poorly metastatic cells about 3-fold, whereas the secretome from cells cultured on stiffer substrates (47 kPa) increased the migration speed about 1.5-fold.

**Figure 4:**
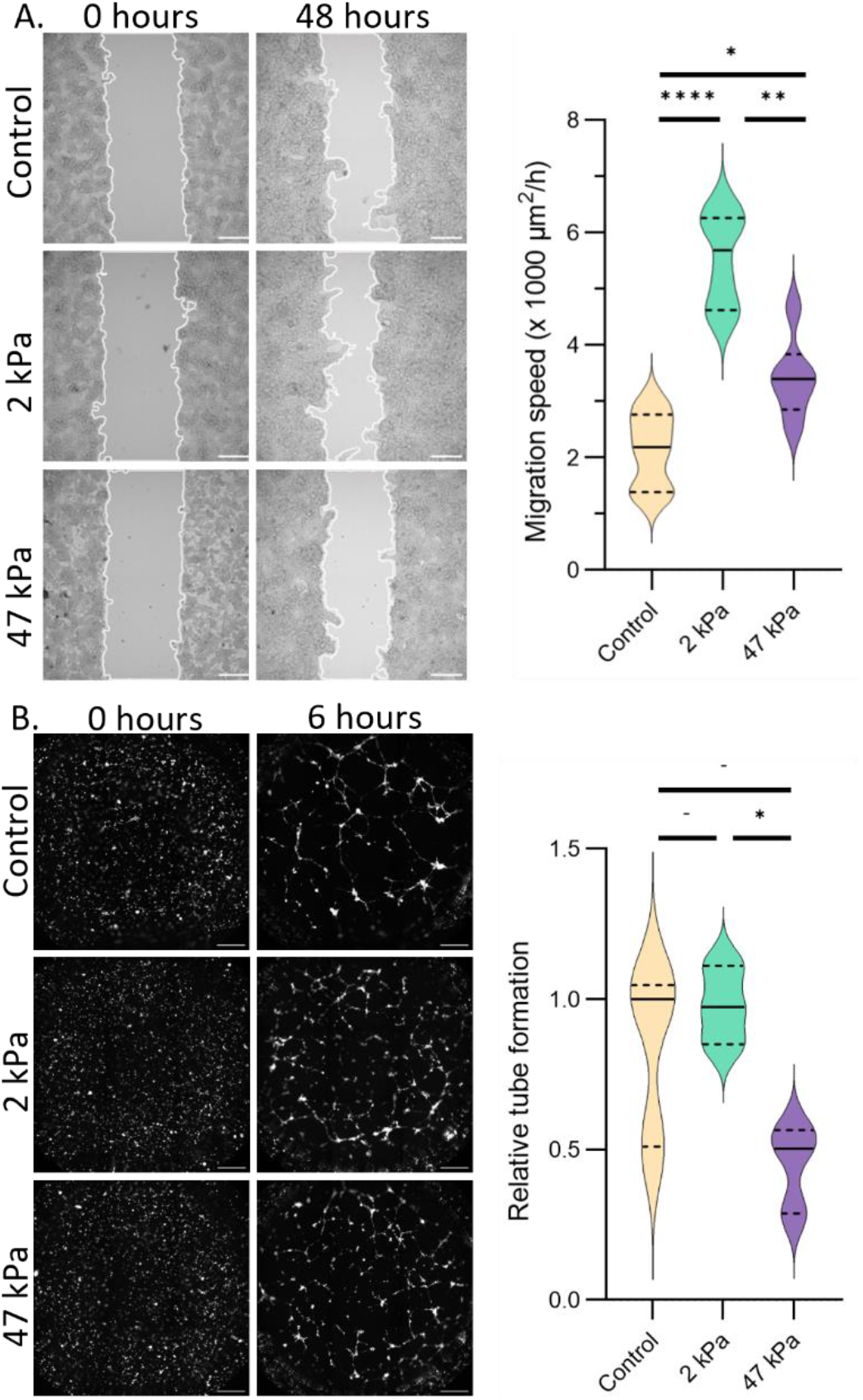
Analyzing functional effects of secretome on cell migration (A) and angiogenesis (B). **A:** Cell migration assay for KM12C wound healing assays incubated with control (lyophilized starvation medium), softer (2 kPa) and stiffer (47 kPa) lyophilized secretomes at different timepoints. Representative images with 200 µm scale bars are shown. Quantified data was obtained from minimally two different replicate samples, from three regions per sample. **B:** Angiogenesis tube formation assays for HUVEC tube formation incubated with control (positive control with complete medium showing maximum tube formation potential), softer (2 kPa) and stiffer (47 kPa) lyophilized secretomes at different timepoints. Representative images with 500 µm scale bars. Quantified data shows the increment of the normalized cell area through time that was normalized to the median of the control condition. Data was obtained from three different replicate samples. **A**,**B:** Data is represented in a violin plot where solid lines show medians, while dashed lines show quartiles. Significance levels: (-) p > 0.05, non-significant; (^*^) p ≤ 0.05; (^**^) p ≤ 0.01; (^****^) p ≤ 0.0001.

#### 4.2 Effects on angiogenesis

Efficient supply of nutrients and removal of waste products is essential for the outgrowth of solid tumors. Given the dysregulation of factors associated to blood vessel formation and remodeling, we evaluated the effect of secretome on the formation of novel blood vessels. Angiogenesis assays revealed that the secretome from KM12L4a cells cultured on softer substrates (2 kPa) was nearly twice as effective in inducing angiogenesis compared to that from cells grown on stiffer substrates (47 kPa), approaching the maximum angiogenic potential that can be observed in this assay (positive control with complete medium) (Fig. 4B).

#### 4.3 Effects on ECM production and degradation

The growth of solid tumors is accompanied by a continuously evolving ECM, involving the production, remodeling, and degradation of ECM proteins. To explore how the secretome influences these dynamical ECM changes, we analyzed the expression and distribution of three collagens (COL2A1, COL5, and COL6A3), which, according to our proteomic analysis, were overexpressed in the secretome of cells grown on the softer substrate (Fig. 5A and S7). While the intracellular distribution of collagen 5 and 6A3 was similar between softer and stiffer substrates, collagen 2A1 exhibit a higher intracellular signal on stiffer substrates. This may be due to reduced secretion of collagen 2A1 on stiffer substrates, resulting in intracellular accumulation. The expected increase in secreted collagen on softer substrates was not observed, possibly due to the removal of loosely attached extracellular collagen during the immunolabeling process.

**Figure 5:**
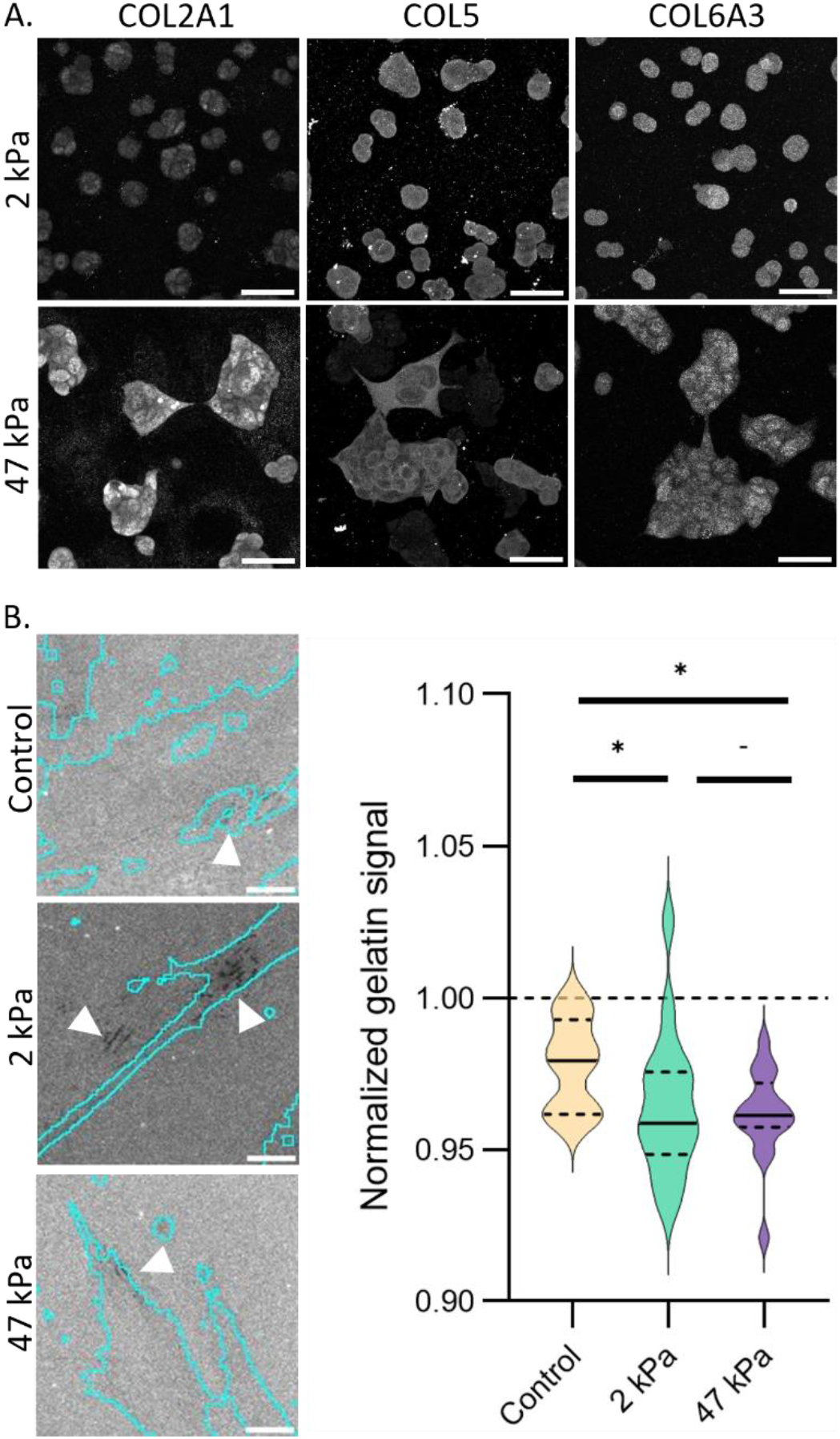
Analyzing functional effects of secretome on ECM production and degradation for stiffer (47 kPa) versus softer (2 kPa) conditions. **A:** Collagen protein expression. Representative images for the maximum z-projection of immunolabeled collagens for KM12L4a cells grown on top of softer (2 kPa) and stiffer (47 kPa) substrates. Side-by-side images were taken with the same acquisition and display settings. Figure shows representative images of three biological replicates, with three fields of view imaged per replicate. The full dataset is available in Fig. S7. Scale bars: 50 µm. **B:** Gelatin degradation assay. Representative images of CAF cells (blue lines show segmented cell boundary) incubated with different lyophilized secretomes to assess the effect on gelatin degradation. Arrows indicate darker regions where gelatin was degraded. Scale bars 20 µm. Violin plot shows quantification of the normalized gelatin signal (ratio of intracellular *versus* extracellular gelatin signal per field of view), for data obtained from minimally 8 different samples from minimally 2 different replicates. Data is represented in a violin plot where solid lines show medians, while dashed lines show quartiles. Significance levels: (-) p > 0.05, non-significant; (^*^) p ≤ 0.05; (^**^) p ≤ 0.01; (^****^) p ≤ 0.0001.

In addition, we evaluated the effect of secretome on ECM degradation, a process often mediated by colorectal cancer-associated fibroblasts (CAFs) in the tumor microenvironment. Using fluorescently labeled gelatin-coated coverslips, we assessed the effect of the secretome on gelatin degradation by CAFs (Fig. 5B). Secretomes from different stiffness conditions showed no significant differences, possibly because the cells produced their own ECM over time, which may have masked any secretome-driven changes. However, secretomes harvested from both softer and stiffer conditions differed significantly from the control (lyophilized starvation medium), suggesting a potential role for the secretome in regulating matrix dynamics.

Collectively, these results confirm that the dysregulation of the secretome due to the stiffness of the surrounding tissue would have a profound effect on the behavior of CRC cells *in vivo*, affecting tumor progression in CRC patients.

## Discussion

In this study, we explore how ECM mechanical properties influence CRC, focusing on protein expression changes in response to the stiffer environments observed during tumor progression. Our findings show that the highly metastatic KM12L4a cells are mechanosensitive, with their morphology changing with substrate stiffness, making them suitable for studying the impact of altered ECM stiffness.

Despite changes in cell morphology, our TMT-based proteomic analysis reveals that substrate stiffnesses above 2 kPa do not alter global intracellular protein expression in KM12L4a cells. Other proteomic studies on liver cancer and endothelial cells reported significant stiffness-related effects, with 240-350 dysregulated intracellular proteins (for a similar total protein count), but direct comparison is difficult due to differences in cell lines and stiffnesses used.^17,19^ In CRC, other factors have been shown to impact intracellular protein expression. For example, 566 proteins were dysregulated when comparing adenoma and adenocarcinoma tissue samples from CRC patients to healthy tissue, reflecting the effect of CRC progression, and 1318 proteins were dysregulated in the KM12C model, indicating the influence of the metastatic potential.^29,31,60^ Although no significant stiffness-induced changes were observed in intracellular proteins, the reliability of our analysis is confirmed by the high number of proteins quantified, and the dysregulation detected in the parallel secretome analysis. Importantly, a lack of dysregulation does not mean intracellular proteins remain unaffected; changes in protein organization or subcellular localization would not be detected in our analysis. In fact, previous spatial proteomic studies have demonstrated that many dysregulated proteins shift in localization, and our tubulin analysis shows unchanged global expression levels despite evident changes in cell morphology and cytoskeletal structure.^29,31^

Unlike intracellular proteins, secreted proteins are significantly influenced by substrate stiffness, as shown by LC-MS/MS and cytokine analyses, and further validated for most selected proteins using dot blot and ELISA tests.

*In silico* analyses confirm that 28% of the dysregulated secretome proteins are released via both classical and non-classical secretion pathways. Notably, 46% of these proteins have been linked to exosomes, with 17% specifically associated with CRC. Exosomes, small extracellular vesicles (40-160 nm), transport bioactive cargo that can influence the ECM and surrounding cells.^61–63^ Previous reports show that substrate stiffness affects exosome composition, with stiffer substrates inducing an increase in molecules involved in cell-ECM interactions (e.g. thrombospondin 1, integrins), ECM degradation (e.g. MMP9), and inflammation and immune evasion (e.g. S100-family).^18,20,21,64,65^ These proteins classes align with our GO and Reactome analyses, which indicate that dysregulated secretome proteins are involved in cell migration and adhesion, intracellular transport, cell death and proliferation, and protein and RNA modifications.

Importantly, substrate stiffness appears to have a more pronounced effect on the secretome than metastatic potential. In our analysis, 44% of the identified proteins were dysregulated due to changes in stiffness, while only 6-12% were linked to metastatic potential in previous studies, including the KM12 model. ^49,59^ Although direct comparisons are challenging due to differences in cell lines and methodologies, the stark difference in the percentage of dysregulated proteins highlights the dominant critical role of stiffness. Additionally, the overlap in affected proteins suggests that both stiffness and metastatic potential may act together to regulate similar biological processes, potentially promoting metastasis. This is consistent with models in other cancer types, such as breast cancer, where oncogenic mutations enhance cell sensitivity to matrix stiffness, reinforcing the interplay between these factors.^20^

Analysis of the effect of the different secretomes on cell behavior shows that the secretome from metastatic cells grown on softer substrates increases the migration speed of poorly metastatic CRC cells and induces angiogenesis more effectively than secretome from cells grown on stiffer substrates. This suggests that metastasized cells, while forming metastases in relatively soft secondary organs, may enhance the migration of poorly metastatic cells at the primary tumor. The interplay between primary tumors and metastases – including micro-metastasis dormancy and reactivation, and metastatic reseeding – is increasingly recognized as integral to tumor progression and supports our hypothesis.^66–68^ Additionally, our angiogenesis assays suggest that metastasized cells can rapidly develop vasculature in newly colonized sites, potentially critical for supporting the rapid growth of metastases.

Our bioinformatic analyses also identified proteins associated with the extracellular matrix, prompting us to evaluate how substrate stiffness impacts collagen deposition and gelatin. Although proteomic data indicated increased ECM dynamics on softer substrates, our functional assays showed no differences (except for intracellular distribution of COL2A1). While we adhere to standardized, extended timescales to replicate *in vivo* scenarios, future research could focus on the impact of self-deposited ECM to better assess the effect of the secretome. Despite finding no clear impact of stiffness, our analyses indicate that matrix deposition and degradation are stimulated by the secretome, regardless of matrix stiffness. This suggests a positive feedback loop where substrate stiffness and remodeling gradually increase during tumor progression (as previously suggested).^6,8^

Our findings underscore the pivotal role of stiffness-induced, secretome-mediated stiffness-induced changes in CRC development and progression. This study is the first to examine the effects of the entire secretome, revealing its broader impact beyond specific secreted fractions such as exosomes, and emphasizing its key role in cancer progression. We expect these findings to form a strong basis for future research into secretome-ECM interactions and inspire the development of more targeted therapies.

## Author contributions

C.C. and S.A. optimized the workflow for the synthesis and functionalization of PAAm hydrogels, under the supervision of S.R. Rheological characterization was performed by C.C., with guidance from B.S. and L.G., and under supervision of P.K. LC-MS/MS experiments and data analysis were performed by A.M.-C. with assistance of C.C. and under the supervision of R.B. Validation experiments, functional assays, and data analysis were performed by C.C., A.M.-C. and G.S.-F. under supervision of R.B. and S.R. The manuscript was drafted by C.C. and R.B., with input from A.M.-C. and G.S.-F., and corrections by S.R. Experiments were conceptualized, designed, and interpreted by C.C., A.M.-C., G.S.-F., R.B., and S.R., and the final manuscript was written with input from all authors.

## Acknowledgements

We thank Fidler’s lab (MD Anderson Cancer Center) for sharing KM12 model cell lines, De Wever’s lab (UGent) for sharing the CAF CT5.3 cell line, and Prof. E. Planus for sharing details regarding the gelatin degradation protocol. Support of the Advanced Optic Microscopy Unit of the Instituto de Salud Carlos III is gratefully acknowledged.

This work was funded by financial support of PID2022-140307OB-I00 funded by MCIN/AEI/10.13039/501100011033 and by “ERDF A way of making Europe” to R.B., PI20CIII/00019 and PI23CIII/00027 grants from the AES-ISCIII program, co-financed with FEDER funds, to R.B. This project has received funding to R.B. from the EIC Pathfinder program 2023 of the European Innovation Council (EIC) under Grant Agreement #101130574. Additionally, S.R. gratefully acknowledges KU Leuven financial support through internal funds (IDN/20/021, KA/20/026, C14/22/085). This work was also funded by the Research Foundation Flanders (FWO) through a project with grant number G0C2422N. In addition, C.C. and S.A. are recipients of an FWO PhD fellowship for fundamental research, grant numbers 1121223N and 1S95123N. G.S.-F. is recipient of an FWO junior postdoctoral fellowship, grant number 12AML24N. C.C. received an FWO travel grant for a long stay abroad.

