## Supplemental information_figures and table legends for "Deciphering stiffness-driven changes in colorectal cancer by proteomics"

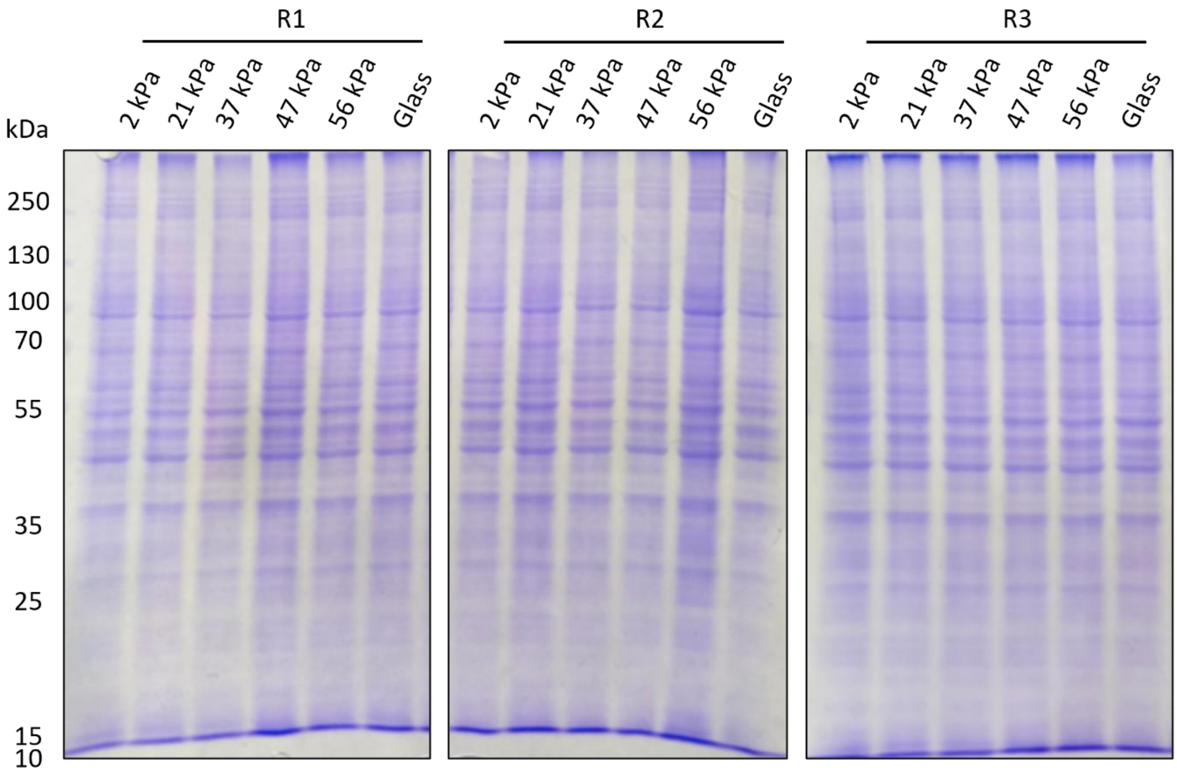


**Figure S1**: **Cell protein extracts visualization to ensure sample quality prior to TMT-labeling.** Samples were visualized on a 10% SDS-PAGE gel under reducing conditions and Coomassie blue staining. Note that 47 kPa_R1 and 56 kPa_R2 were observed to be 12.5% denser and corrected prior to TMT-labeling. "R" denotes "replicate".

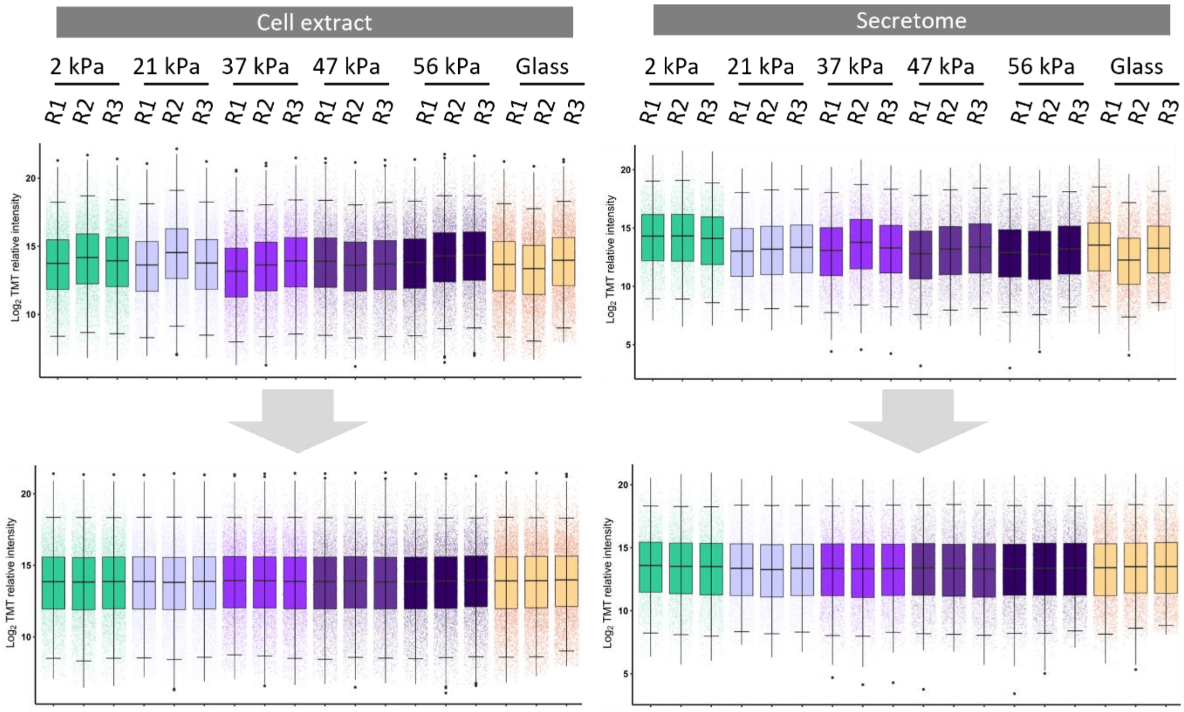


**Figure S2: Normalization of raw data from MaxQuant.** The box plots represent the Log2 of the median protein intensities of cell extracts (left) and secretomes (right) before (top) and after (bottom) sample loading (SL) normalization. "R" denotes "replicate".

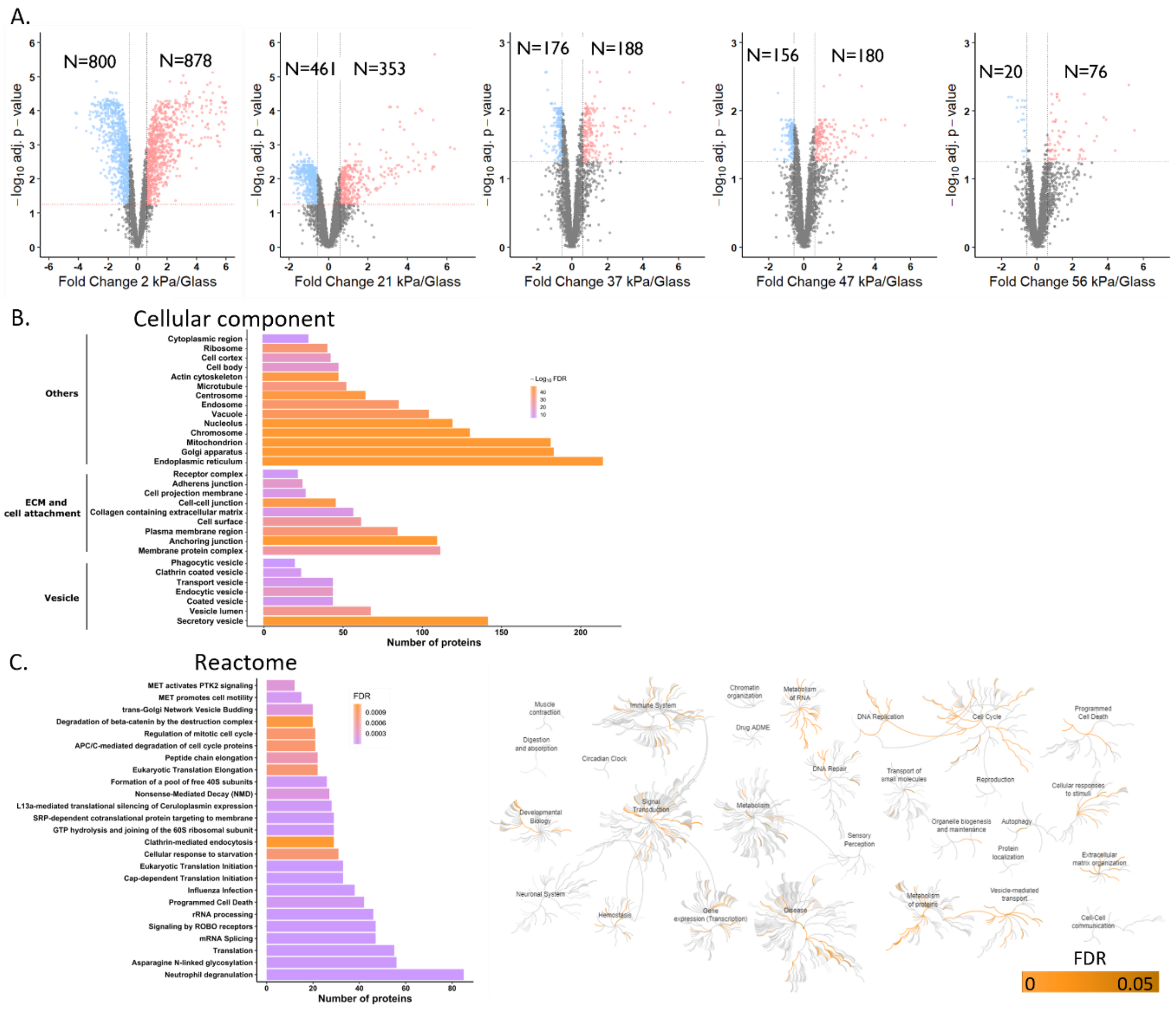


**Figure S3: Additional info for proteomic analyses of secretome samples. A:** Volcano plots of proteins identified and quantified as altered in PAAm hydrogels (2, 21, 37, 47, or 56 kPa) in comparison to glass conditions in secretomes. The x-axis represents Log2 expression ratio (fold change) of hydrogels *versus* glass relative protein expression ratios, the y-axis depicts the FDR value (adj. p-value) based on -Log10. Colored dots represent differentially expressed proteins upregulated (red) and downregulated (blue) in stiff conditions with FDR ≤ 0.05 (represented by red dashed horizontal line) and 1.5-fold expression ratio (represented by two black dashed vertical lines). **B-C:** Bioinformatics analysis of dysregulated proteins in stiff (21, 37, 47, and 56 kPa) *versus* soft (2 kPa) conditions. **B:** Bar plots represent the most significantly enriched cellular components in which dysregulated proteins are involved according to Gene Ontology (GO) analysis. Color gradient from purple to orange represents the -Log10 FDR. **C**: Reactome analysis (FDR < 0.05) revealed the most significantly altered pathways due to protein dysregulation in stiff conditions. Significance is shown with the FDR in a purple to orange color gradient (bar plot) or orange gradient (nodes).

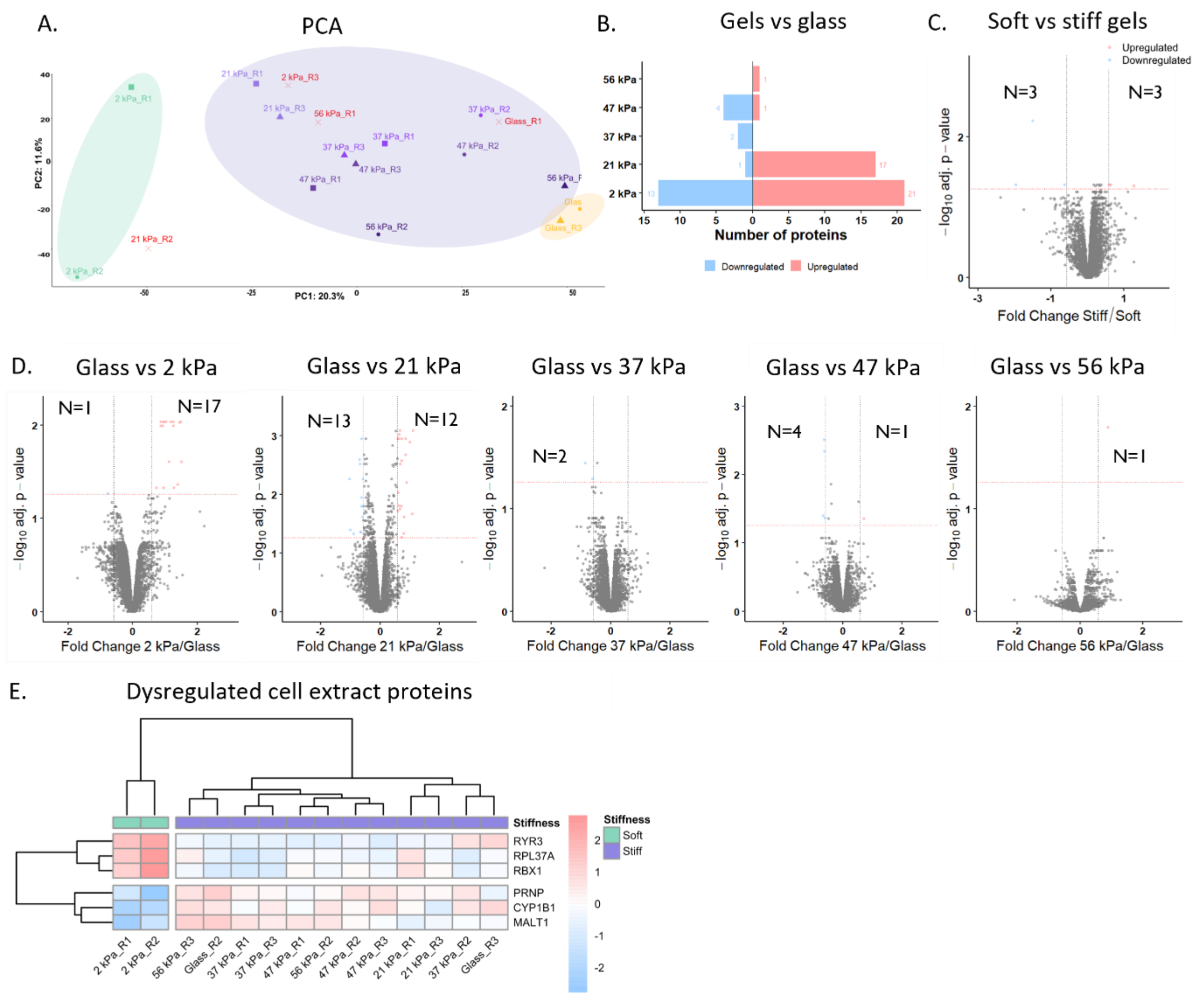


**Figure S4: Proteomic analyses of cell extracts. A-C:** Differential expression analyses. **A:** Principal component analyses (PCA) representing sample clustering according to the stiffness. **B:** Bar plots with the number of upregulated (red) and downregulated (blue) proteins in the different stiffness conditions, compared to glass. Volcano plots with the fold changes and p-values are shown in panel D. **C:** Volcano plots of proteins identified and quantified as altered in stiff (21, 37, 47, and 56 kPa) *versus* soft (2 kPa) conditions. The x-axis represents the $Log_2$ expression ratio (fold change) of stiff *versus* soft relative protein expression ratios, the y-axis depicts the FDR value (adj. p-value) based on -Log10. Colored dots represent differentially expressed proteins upregulated (red) and downregulated (blue) in stiff conditions with FDR ≤ 0.05 (represented by red dashed horizontal line) and 1.5-fold expression ratio (represented by two black dashed vertical lines). **D:** Volcano plots of proteins identified and quantified as altered in PAAm hydrogels (2, 21, 37, 47, or 56 kPa) in comparison to glass conditions. The x-axis represents Log2 expression ratio (fold change) of hydrogels *versus* glass relative protein expression ratios, the y-axis depicts the FDR value (adj. p-value) based on -Log10. Colored dots represent differentially expressed proteins upregulated (red) and downregulated (blue) in stiff conditions with FDR ≤ 0.05 (represented by red dashed horizontal line) and 1.5-fold expression ratio (represented by two black dashed vertical lines). **E:** Bioinformatics analysis of dysregulated proteins in stiff (21, 37, 47, and 56 kPa) *versus* soft (2 kPa) conditions. Heatmap of the TMT relative protein expression of the six dysregulated proteins identified in the cell extracts TMT analysis normalized to the mean row intensity.

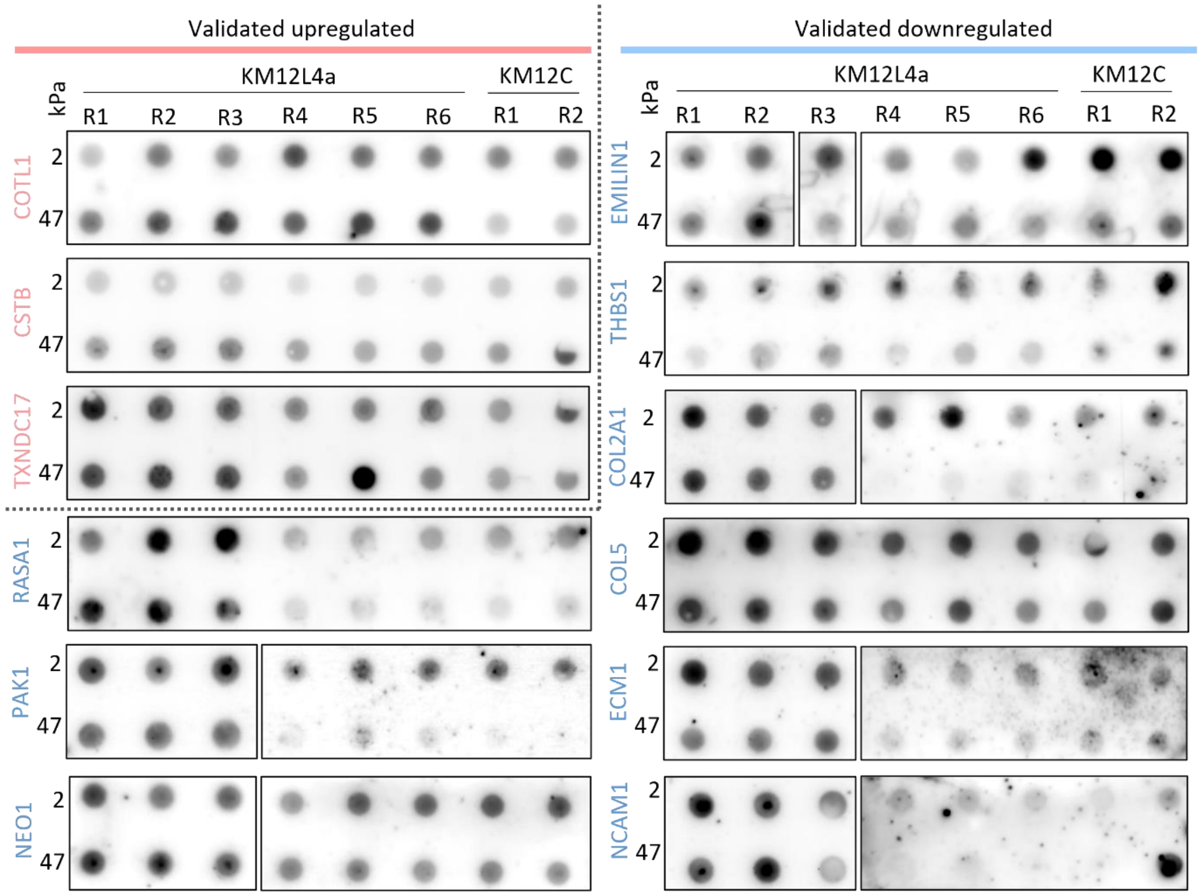


**Figure S5: Dot blot analyses for selected upregulated and downregulated secretome proteins quantified in Fig. 3A.** White intersections within one membrane indicate representations taken from different signal development reagents or acquisition settings. Brightness and contrast of images are adjusted equally per membrane section without omitting signal, to better represent the observed differences. For EMILIN1, KM12L4a 2 kPa_R3 and 47 kPa_R3 samples were accidently swapped during sample loading, hence the correction in the figure. For COL2A1, results for KM12L4a R4-6 and KM12C R1-2 were obtained from 25 µL instead of 5 µL non-lyophilized secretome. “R” denotes “replicate”.

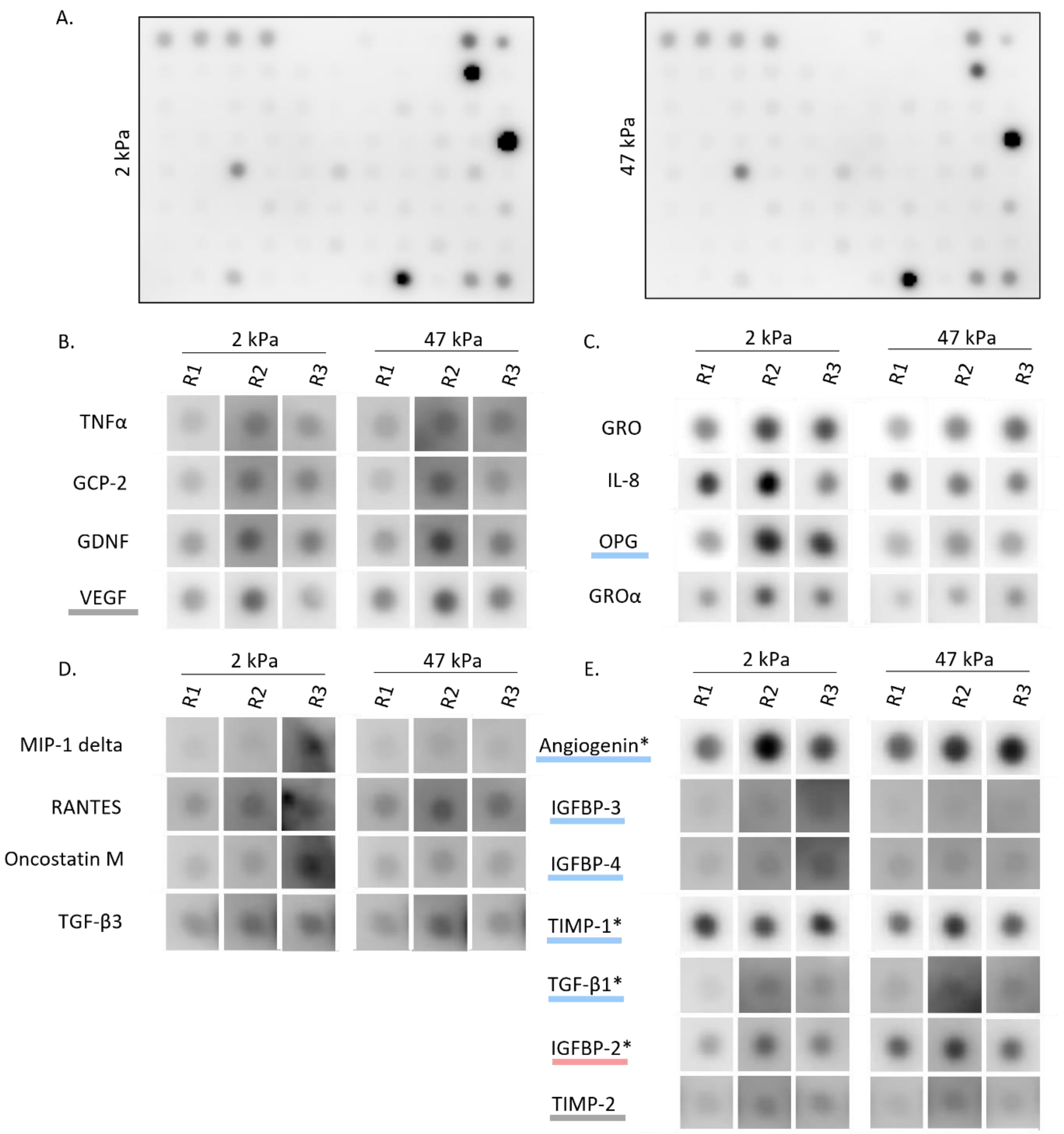


**Figure S6: Cytokine arrays for stiffer (47 kPa) versus softer (2 kPa) KM12L4a original secretomes (R1-3)**. **A:** Representative images of 80 cytokines analyzed using the C5 human cytokine array from RayBiotech. **B-C:** Upregulated (B) or downregulated (C) cytokines as quantified in Fig. 3B. For meaning of colored lines, see panel E. **D:** Cytokines excluded due to low signal intensity or signal overlap with neighboring spots. **E:** Cytokines also identified in LC-MS/MS analyses (underlined), although they are non-significant in this cytokine array. Cytokines can be close to equal levels (grey line), upregulated (red line), or downregulated (blue line). Note that VEGF and OPG were also identified in LC-MS/MS analyses (underlined in panels B-C). **B-E:** To better represent the observed differences, acquisition settings, brightness and contrast were adjusted equally per cytokine without omitting signal. However, settings may differ between different cytokines.

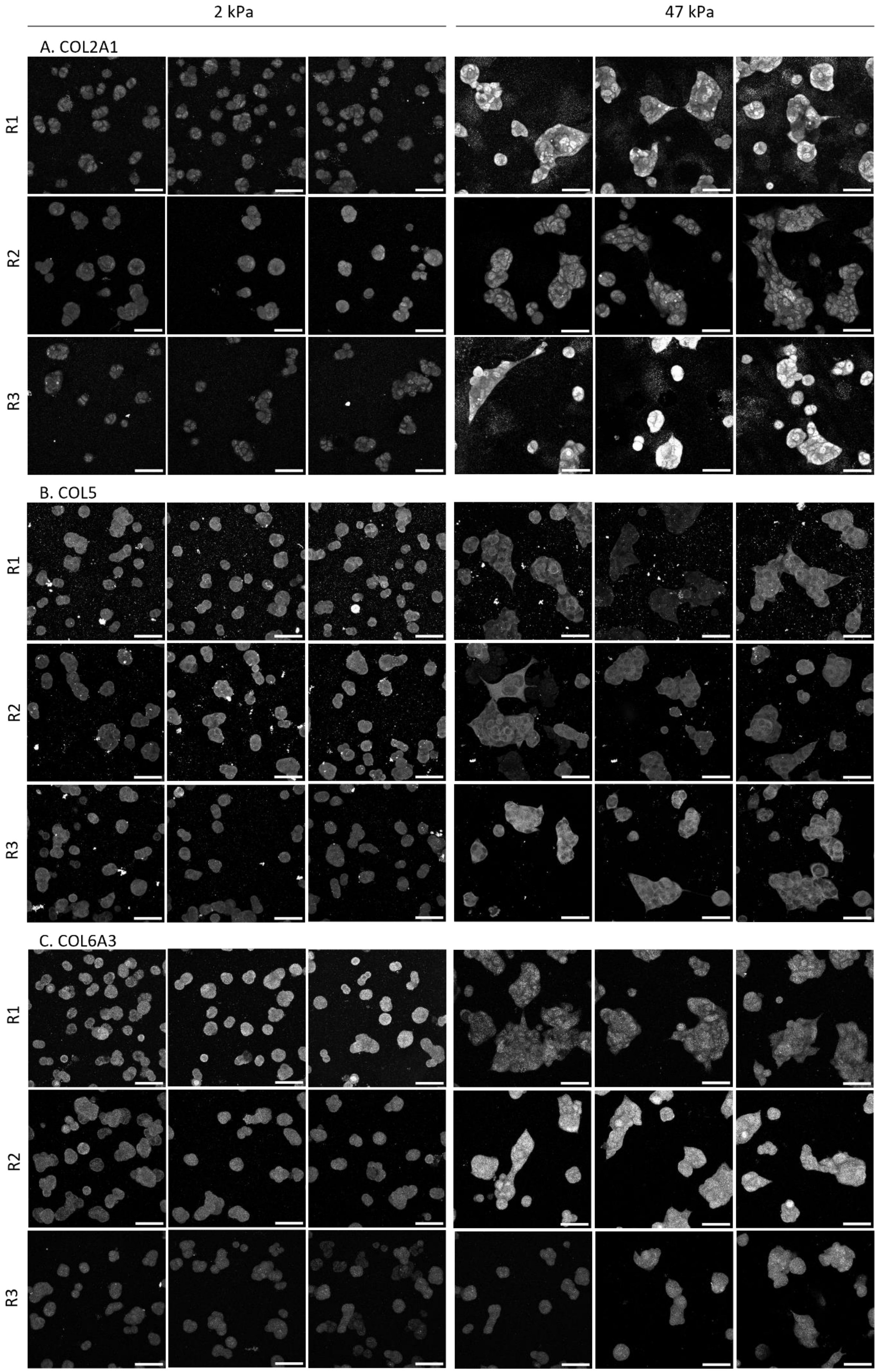
2 kPa
47 kPa
A. COL2A1
B. COL5
C. COL6A3
R1
R2
R3

**Figure S7: Full dataset for collagen protein expression.** Maximum z-projection of different collagens in KM12L4a cells grown on softer (2 kPa) and stiffer (47 kPa) substrates. Side-by-side images were taken with the same acquisition and display settings. Figure shows full dataset of images represented in Fig. 4C, with three fields of view per replicate. Note that these replicates are unrelated to the replicates used in proteomics. “R” denotes “replicate”. Scale bars 50 µm.

**Table S1 (see excel document): Details for PAAm hydrogel stiffnesses.** Practical gel composition denotes quantities (in µL) necessary for production of one large gel (total volume 10 mL, 9.5 mL necessary for gel synthesis). Detailed stiffness characterization shows rheology parameters and results. Initial gap and axial force are read out upon uniform contact between the geometry and the PAAm gel, and a non-negligible axial force of minimally 0.01 N. Final gap and axial force are read out before zeroing of the axial force and starting the measurement.

**Table S2 (see excel document): Reagents employed for labeling of targets.** For blocking buffers, “FBS” means 10% FBS in DPBS, “milk” means 3% skimmed milk and 0.1% Tween 20 in DPBS, “BSA” means 3% BSA powder and 0.1% Tween 20 in DPBS.

**Table S3 (see excel document): List of proteins identified in the secretome TMT 18-plex analysis.** The expression ratio, FDR (adjusted p-value), and dysregulation observed are shown for the most interesting differential expression analysis performed.

**Table S4 (see excel document): List of proteins identified in the cell extract TMT 18-plex analysis.** The expression ratio, FDR (adjusted p-value), and dysregulation observed are shown for the most interesting differential expression analysis performed.

**Table S5 (see excel document): Analysis of secretory pathways of the 1476 dysregulated proteins between 2 and 47 kPa secretomes.**
